# Gene-level complexity explains genome-wide variation in the distribution of fitness effects

**DOI:** 10.64898/2026.04.08.717178

**Authors:** Burçin Yıldırım, Jennifer E. James

## Abstract

The distribution of fitness effects (DFE) — describing how harmful, neutral, or beneficial new mutations are — is central to understanding how populations evolve. Although the DFE varies across genomes and species, it remains unclear which aspects of genomic organization drive this variation. Here, we inferred gene-level selective constraints across the genomes of *Mus musculus castaneus*, *Drosophila melanogaster* and *Saccharomyces cerevisiae* using a combination of population genetics and machine learning trained on diverse gene features. Many gene features were predictive of selective constraint, with conservation, gene structure, and expression being the most informative. These selective constraints delineated gene classes with distinct DFEs. Genes with higher connectivity and expression — features reflecting how many traits a gene influences — experienced stronger and less dispersed deleterious effects with increasing selective constraint. Between species, the rate of adaptation decreased with increasing organismal complexity, whereas across the genome it did not decrease monotonically with selective constraint, but tended to be higher at intermediate levels. While between-species comparisons of DFE parameters were less consistent with predictions of Fisher’s geometric model (FGM) based on organismal complexity, variation in DFE parameters across the genome aligned more closely with FGM when complexity was considered at the gene level. Our results suggest that gene-level complexity, captured by genomic feature proxies, provides a more informative definition of complexity for DFE variation than organism-level labels, and highlight the value of using gene features collectively to link genomic architecture, fitness landscapes, and patterns of molecular evolution.

## Introduction

Mutations arise in all organisms, affecting phenotypes and, consequently, fitness. Their fitness consequences are typically described by the distribution of fitness effects (DFE), which spans a continuum from deleterious through neutral to beneficial mutations (Eyre-Walker and Keightley, 2007). The fitness effect of a mutation influences the frequency at which it segregates in a population, shaping the evolutionary fate of populations, relating to questions in evolutionary and quantitative genetics, such as the amount and nature of genetic variation, the rate of adaptation, and the genetic architecture of traits. Understanding the DFE is therefore central to characterizing many evolutionary process.

Considerable effort has been devoted to characterizing the DFE (Eyre-Walker et al., 2006; Keightley and Eyre-Walker, 2007; Tataru et al., 2017) and its variation between species and across the genome. Between closely related species the DFE is relatively stable, and correlations between DFEs decrease with genetic differentiation (Castellano et al., 2019; Huang et al., 2021; Lin et al., 2025). The DFE is often similar among populations of the same species, and even between domesticated species and their wild relatives despite distinct demographic histories (Chen et al., 2017; James et al., 2023; Amorim et al., 2024). These results suggest that the DFE is not highly sensitive to changes in demography or environment, but is instead more constrained by species biology — for example, life-history traits and genome characteristics — which tend to be similar among closely related species.

Consistent with the importance of long-term processes, several studies have found relationships between life-history traits and the DFE (Chen et al., 2017; Muyle et al., 2021). In addition, analyses of genomic properties have begun to uncover their influence on the DFE. Bergman and Eyre-Walker (2019) and Moutinho et al. (2019) have shown that knowledge of amino acid properties and protein structure, respectively, improves our understanding of DFE characteristics. Similarly, the analyses of Hämälä and Tiffin (2020) on the factors affecting selective constraints revealed that gene expression level and gene network connectivity are strong predictors, and Chen et al. (2022) found that the level of sequence conservation covaries with the DFE parameters. Finally, Moutinho et al. (2022) found that gene age was negatively related to the rate of adaptive and nonadaptive evolution, which they interpreted as evidence that young genes are under weaker selective constraint. While these studies identified some genomic predictors, we still lack an understanding of which other features contribute to DFE variation at the genomic level, the extent of this variation, and a more systematic description of it..

Theoretically, Fisher’s geometric model (FGM) is one of the most commonly used frameworks to predict DFE characteristics and variation (Fisher, 1930; Martin and Lenormand, 2006; Lourenço et al., 2011; Tenaillon, 2014). Originally introduced as a model of adaptation describing the probability of mutations being advantageous, it also provides a basis for predicting the full distribution of fitness effects using a small set of assumptions. In FGM, an organism is represented as a vector of *n* traits determining its position in an *n*-dimensional phenotypic space, and fitness depends on the distance from this position to a given optimum. Mutations displace phenotypes in this space, and the resulting DFE can be derived from these displacements. The fitness effect of a mutation—whether beneficial or deleterious, large or small—depends on whether the mutation moves the phenotype closer to or farther from the optimum (Orr, 1998). Therefore, this geometric mapping to an optimum is a core component of the model that affects the DFE.

Another core concept in FGM is complexity, although this is difficult to define precisely. Theoretically, complexity is described by the number of independent traits under selection, that is, by the dimension of the phenotypic space, *n*. The classic formulation of Fisher’s geometric model assumes universal pleiotropy, where every mutation affects all traits in an organism; under this assumption, pleiotropy and complexity effectively overlap. In this case, *n* is determined by the overall dimensionality of the phenotypic space, and the focus is therefore on organismal complexity (Tenaillon et al., 2007; Tenaillon, 2014). As the dimensionality increases, the probability that a mutation is beneficial decreases, a phenomenon known as the cost of complexity (Orr, 2000). Consequently, new mutations are more likely to be deleterious in complex organisms than in less complex ones, potentially leading to relative differences in their DFEs. Furthermore, complexity constrains the rate of adaptation by increasing the likelihood that alleles beneficial for some traits have deleterious effects on others. As a result, adaptation should mainly proceed through the fixation of small-effect mutations (Orr, 2005).

To date, only a few studies have corroborated the expectations of FGM under universal pleiotropy, showing a higher proportion of deleterious mutations in humans compared to *Drosophila* (Huber et al., 2017), or suggesting that, on average, mutations are more deleterious in mammals than in birds and insects (Lin et al., 2025). To our knowledge, the relationship between the rate of adaptation and proportion of beneficial mutations and organismal complexity has never been tested directly, although some previous results were broadly consistent with FGM expectations — for example, the lower adaptive substitution rates in mammals compared to insects (Rousselle et al., 2020) and the higher proportion of beneficial mutations in humans compared to mice and *Drosophila* (Zhen et al., 2021). Inconsistencies or the lack of a fine-scale characterization may arise when the classic assumptions of FGM are violated.

One key assumption of the FGM that has received much scrutiny is that of universal pleiotropy or isotropic mutational effects (i.e., equal and independent effects across all traits). Mutational pleiotropy can be modular rather than universal, with most mutations directly influencing only traits within their developmental module (Wagner and Zhang, 2011). More broadly, development structures the genotype-phenotype map such that mutational effects tend to be aligned along developmental and regulatory axes rather than distributed isotropically across all phenotypic dimensions (Salazar-Ciudad, 2021; Rohner and Berger, 2025). Consequently, even highly pleiotropic mutations can have low effective dimensionality if their effects on different traits are correlated — a point that holds even under universal pleiotropy (Hill and Zhang, 2012). Similarly, other alternative assumptions regarding pleiotropy that predict variable degrees of pleiotropy across mutations or correlated effects — such as restricted pleiotropy, where mutations affect only a subset of traits rather than all traits, and synergistic pleiotropy, where effects on multiple traits are aligned in the same direction — would result in reduced effective dimensionality (Wang et al., 2010). Altogether, these considerations shift the focus from organismal complexity, defined by the total dimension of the phenotypic space, to gene-level complexity, defined by the effective dimensionality of mutations at a given gene (Walsh and Lynch, 2018). Such gene-level complexity could in principle better explain variation in the DFE both across the genome and between species.

Indeed, some studies have reported that the observed degree of pleiotropy across mutations is not equal but follows an L-shaped distribution, with most mutations affecting only a few traits (Wagner et al., 2008; Wang et al., 2010; Wagner and Zhang, 2011). It has been shown that highly pleiotropic genes are associated with higher expression and more protein-protein interactions (He and Zhang, 2006; Barbitoff et al., 2025), such that these features can serve as proxies for pleiotropy, and therefore gene-level complexity. More importantly, these studies proposed that modular pleiotropy can maximize the rate of adaptation in organisms with intermediate levels of complexity (Wagner et al., 2008; Wang et al., 2010). Similarly, Lourenço et al. (2011) developed an extension of FGM, incorporating modular pleiotropy and showing that the rate of adaptation has a concave, rather than a strictly decreasing, relationship with increasing pleiotropy. Martin (2014) further demonstrated that FGM can be derived from the modular structure of phenotypic networks and can better explain variation in the DFE, especially across the genome. Lastly, experimental evolution and local adaptation studies have shown that highly pleiotropic mutations, typically associated with larger effect sizes (Wagner et al., 2008; Wang et al., 2010), can promote rather than impede adaptation (Hämälä et al., 2020; Thorhölludottir et al., 2023; Whiting et al., 2024; Koch et al., 2025), which has been explained by aligned correlated responses among genes within the same regulatory networks.

One way to better understand variation in the DFE therefore seems to be to examine how it relates to genomic features, and whether this variation reflects underlying differences in pleiotropy and complexity across genes, as suggested by FGM and its extensions. However, testing this requires characterizing the DFE across genes, which is methodologically challenging: individual genes have too few polymorphic sites to allow reliable DFE inference. One approach is to group genes that are assumed to share similar DFEs, using genomic features as proxies. Previous studies have taken this approach by grouping genes according to individual features such as expression level, conservation or recombination rate to infer the DFE or related statistics (Castellano et al., 2018; Hämälä and Tiffin, 2020; Chen et al., 2022), providing valuable but necessarily partial insights. Here, we take a different approach by first estimating per-gene selective constraints that incorporate information about their genomic features, and then using these constraint estimates to group genes for downstream DFE inference (Figure 1). The per-gene constraint estimates represent the overall deleteriousness of a gene and serve as a natural proxy for the underlying DFE — genes with similar constraint values are expected to share similar DFEs.

**Figure 1:**
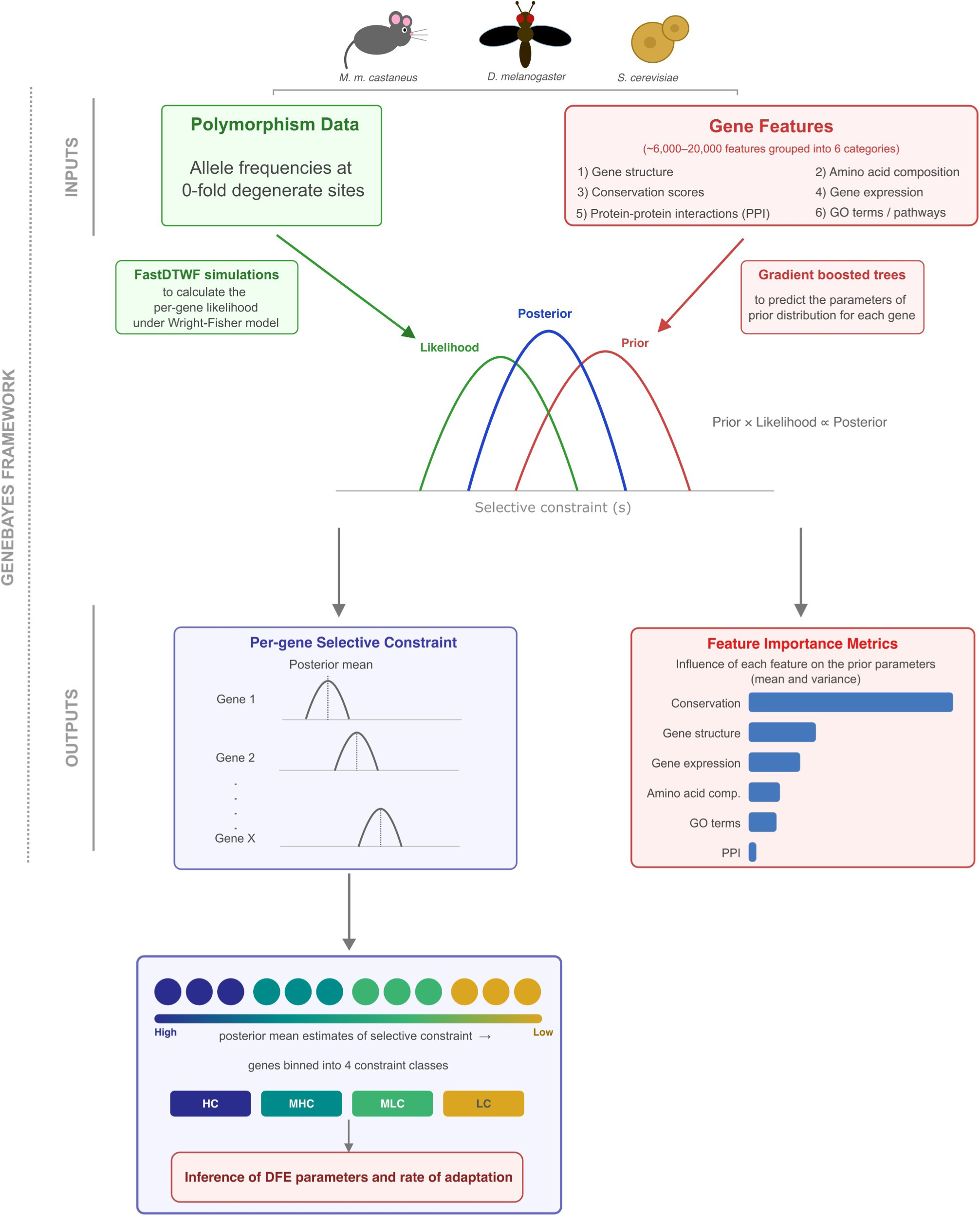
Schematic representation of the GeneBayes framework as implemented in this study. Gene features compiled across six categories (Supplementary Tables 1a–c) were used to learn a prior distribution of selective constraints for each gene via gradient-boosted trees, while simulations under a discrete-time Wright-Fisher model and allele frequencies of 0-fold degenerate variants were used to compute per-gene likelihoods. The first output, feature importance metrics, reflects the contribution of each feature to the learned prior. The second output, posterior per-gene selective constraints, was used to group genes for downstream DFE inference.

Specifically, to estimate gene-level selective constraints, we used GeneBayes (Zeng et al., 2024), an empirical Bayes framework that combines an explicit population genetic model with a machine-learning approach trained on diverse gene features, including conservation scores, gene expression levels, interaction patterns, and gene network properties, across house mice (*Mus musculus castaneus*), fruit fly (*Drosophila melanogaster*), and wild yeast (*Saccharomyces cerevisiae*). Crucially, this framework also allows us to assess how the grouping of genes relates to their genomic properties, linking patterns of DFE variation to gene-level features that can reflect differences in pleiotropy and complexity. With this approach, we ask three main questions: does the DFE vary across the genome, and if so, which gene-level properties are associated with this variation, and can variation in the DFE parameters and the rate of adaptation be systematically explained by theoretical predictions when complexity is considered at the gene level.

## Methods

### Genomic data

We obtained previously published whole-genome data from ancestral-range populations of three species: *Mus musculus castaneus* (mouse) from India (Halligan et al., 2010), *Drosophila melanogaster* (fruit fly) from Zambia (Lack et al., 2015), and wild *Saccharomyces cerevisiae* (yeast) isolates belonging to the Far East Asia clade (Peter et al., 2018). The datasets included 10 diploid, 69 haploid, and 8 diploid genomes for mouse, fruit fly, and yeast, respectively.

For *D. melanogaster*, sequences were obtained as consensus FASTA files, and a full description of data processing is provided in Lack et al. (2015). For mouse, raw sequencing data was downloaded from NCBI (accession number PRJEB2176) and mapped to the mm10 reference genome using bwa-mem with default parameters (Li, 2013). BAM files were generated with samtools (Li et al., 2009) and cleaned using PicardTools (https://broadinstitute.github.io/picard/) after fixing mate information between paired reads. Callable sites were identified, and individual genotype VCF (GVCF) files were generated and jointly genotyped using the Haplo-typeCaller, GenomicsDBImport, and GenotypeGVCFs tools of GATK (McKenna et al., 2010). From the resulting VCF files, we retained only biallelic SNPs and invariant sites. Variant filtering followed these criteria: (i) MQ < 55, (ii) FS > 60 and SOR > 3, (iii) variant confidence (QD) < 4, (iv) MQRankSum < –5 or > 5, (v) ReadPosRankSum < –4 or > 4, and (vi) total read depth (DP) across all samples < 4. Sites with missing data were removed using VCFtools (Danecek et al., 2011). For yeast, we obtained GVCF files mapped to the R64 reference genome from Loegler et al. (2024), and analysis-ready VCF files were produced following the same procedure as for mouse. Because *S. cerevisiae* is a highly selfing species and therefore expected to show excess homozygosity and an overrepresentation of even-frequency alleles (Blischak et al., 2020), we haploidized the yeast genome by converting diploid genotypes to haploid in the VCF file through random sampling of one haplotype per individual to avoid bias in downstream analyses.

Based on the genome annotations available for each reference genome, we extracted protein-coding genes, selecting the longest transcript for genes with alternative isoforms. For downstream analyses, we restricted our data to 0-fold and 4-fold degenerate sites, which were annotated using custom scripts.

### Gene features

For all species, we compiled 6 categories of gene features: (i) gene structure, (ii) amino acid composition, (iii) conservation scores, (iv) gene expression, (v) connectedness in pro-tein–protein interaction (PPI) networks, and (vi) Gene Ontology (GO) terms and/or biological pathways. Individual features within each category are listed in Supplementary Tables 1a–c and in the files provided in the repository (https://github.com/Burciny/DFEvariationAcrossGenomes). Briefly, gene structure features included GC content and length of the CDS, 3’-UTR, 5’-UTR, and transcript, as well as exon and transcript counts, calculated from the genomic data. Recombination rate was assigned to each gene as follows: for mouse, mean cM values were calculated from markers overlapping each gene using the Cox genetic map (Cox et al., 2009); for fruit fly and yeast, cM values were obtained from published estimates in 100 kb and 20 kb windows, respectively, and assigned to genes overlapping each window (Comeron et al., 2012; Liu et al., 2019). Amino acid composition features included the percentage of each of the 20 amino acids, total amino acid count, and percentages of broader physicochemical groups (hydrophilic, hydrophobic, amphipathic, polar, nonpolar, and charged residues), all calculated directly from the genomic data.

For conservation measures, we first downloaded PhastCons scores for all species from the UCSC Genome Browser (Siepel et al., 2005; Haeussler et al., 2019). PhastCons estimates the probability that each site is conserved across a multiple-species alignment, with higher values indicating stronger conservation. We also obtained SIFT scores for each species from the SIFT 4G database (Ng and Henikoff, 2003) (https://sift.bii.a-star.edu.sg/sift4g/public/; last accessed November 30, 2025). SIFT predicts the functional impact of amino acid substitutions based on sequence homology, with lower scores indicating more deleterious substitutions. As both of these scores are given per-site based, we calculated mean, 95th percentile and maximum values for each gene. Finally, we estimated *d_N_/d_S_* ratios using the codeml program in PAML (Yang, 2007), with codon frequencies estimated from the data (codonFreq = 2) and separate *d_N_/d_S_* values for each branch. To perform this, we generated codon-based alignments of protein-coding genes using pal2nal (Suyama et al., 2006) with one or more closely related species: *D. simulans* and *D. yakuba* for *D. melanogaster*, *Rattus norvegicus* for *M. m. castaneus*, and *S. paradoxus* for *S. cerevisiae*.

Gene expression data for mouse, fruit fly, and yeast were obtained from the Mouse Genome Database (MGD) (Baldarelli et al., 2024), FlyBase (Jenkins et al., 2022), and the Expression Atlas (Papatheodorou et al., 2020), respectively, each of which curates gene expression values across multiple independent studies. Protein-protein interaction data for mouse and yeast were obtained from the Alliance of Genome Resources (Bult and Sternberg, 2023), and for fruit fly from FlyBase (Jenkins et al., 2022). From these datasets, we constructed network graphs and calculated connectedness metrics — degree, betweenness, closeness, and eigenvector centrality, and clustering coefficient — using the Python package networkx (Hagberg et al., 2008). GO terms and biological pathway annotations for fruit fly were obtained from FlyBase (Jenkins et al., 2022), and for mouse and yeast from the Gene Ontology Consortium (Gene Ontology Consortium, 2021). Each GO term was encoded as a binary feature indicating whether a gene was annotated with that term. The final dataset included 19828, 10148, and 6101 features for mouse, fruit fly, and yeast, respectively.

### Estimating per-gene selective constraints

We used the previously developed GeneBayes, an empirical Bayes framework for estimating posterior per-gene selective constraints while accounting for the relationship with multiple gene features (Zeng et al., 2024). The approach consisted of two components: first, a prior distribution of selective constraints for each gene, learned via machine learning trained on the gene-level features we compiled. Specifically, the parameters of the prior distribution were predicted using gradient-boosted trees, as implemented in NGBoost (Duan et al., 2020). Second, a likelihood derived from an explicit population genetics model (see below). By combining these two, the posterior distribution of selective constraints for each gene was obtained (Figure 1). All in all, by incorporating gene-level features, the framework leverages the shared biology of similar genes to improve the constraint estimates obtained from the population genetic model.

Selective constraint here refers to the heterozygous selection coefficient against 0-fold degenerate mutations— unlike the original GeneBayes implementation, which was trained on loss-of-function variant frequencies. As stated in the introduction, to investigate genome-wide variation in the DFE, genes need to be grouped such that each group shares a sufficiently similar DFE to allow reliable inference. Posterior per-gene selective constraints estimated by this framework captures the overall deleteriousness of a gene and serve as a good proxy for the underlying DFE. Therefore, they were used to group genes for downstream DFE inference.

In our implementation of GeneBayes, we followed the same prior parameterization as described in the original study. Before fitting the model, we removed features with no variance and features that had no significant Spearman correlation with the ratio of 0-fold to 4-fold variants per gene (*P*_0_*/P*_4_; *P*_4_ serving as the expectation without selection). The features removed during filtering are listed in Supplementary Tables 1a–c. For each species, roughly 75% of genes were used for training, 14% for validation during hyperparameter tuning, and the remaining genes for testing potential overfitting. Model training stopped if the validation loss did not decrease for ten iterations or when the maximum number of iterations (1000) was

reached. Using this approach, we firstly performed a grid search over seven hyperparameters (Supplementary Table 2). We further tuned the hyperparameters with Bayesian optimization implemented in optuna (Akiba et al., 2019), and used the parameter combination that gave the lowest validation loss from grid search as the starting point.

From the trained model with the lowest validation loss, we obtained posterior mean estimates of selective constraint and feature importance metrics for each gene. Feature importance metrics quantify the influence of each feature on the learned prior, as the relative frequency of their contribution to split the data. These metrics were calculated separately for each parameter of the prior, which in our case were the mean and standard deviation of a logit-normal distribution.

### Population genetic likelihood

The likelihoods used in GeneBayes were calculated using FastDTWF simulations (Spence et al., 2023), which implement a standard discrete-time Wright-Fisher population genetic model (Ewens, 2004; Gillespie, 2004). From these simulations, we obtained the probability of observing an allele frequency at present for a given demography, mutation rate and selection coefficient. For each species, these probabilities were computed across a range of selective constraint values (100 log-linearly spaced points between 10*^→^*^8^ [weak selection] and 1 [lethal], plus 0), resulting in a 101 *↑n* matrix (*n* = sample size). These per-variant likelihoods were then combined into gene-level likelihoods using a composite-likelihood approach: the likelihoods for the observed frequencies of all 0-fold variants in our polymorphism data within a gene were multiplied to obtain the gene-level likelihood for that gene. This resulted in a 101× [number of genes] matrix for each species.

Demographic and mutation rate parameters for *M. m. castaneus* population were obtained from Halligan et al. (2010, 2013); Phifer-Rixey et al. (2020). For Far East Asian *S. cerevisiae*, we used estimates from Liu and Zhang (2021) and Raas and Dutheil (2024) for mutation rate and demography, respectively. For Zambian *D. melanogaster*, demographic parameters were taken from Johri et al. (2020); Ragsdale and Gutenkunst (2017) and mutation rate estimates from Keightley et al. (2014).

### DFE Inference

The DFE inference was performed using fastDFE (Sendrowski and Bataillon, 2024), which builds upon the model implemented in polyDFE (Tataru et al., 2017; Tataru and Bataillon, 2019). fastDFE uses likelihood optimization to infer the DFE from the site frequency spectrum (SFS) while accounting for demography and polarization error through nuisance parameters (Eyre-Walker et al., 2006). The method requires SFS data from two site classes—neutral sites, which are used to model demographic and technical distortions of the SFS, and selected sites, which inform the inference of DFE parameters. Given the unfolded SFS, fastDFE can infer both deleterious and beneficial DFEs.

To construct unfolded SFSs, we inferred the ancestral state at each site using est-sfs (Keightley and Jackson, 2018). For this, we generated reference-genome alignments between each focal species and one or two outgroup species (*D. simulans* and *D. yakuba* for *D. melanogaster*; *Rattus norvegicus* for *M. m. castaneus*; and *S. paradoxus* for *S. cerevisiae*). We assumed the Kimura 2-parameter model of base substitution and performed 10 maximum-likelihood searches. The probability of the major allele at a segregating site being ancestral was used to assign ancestral states. Sites lacking outgroup information or with ambiguous probabilities (0.45 *< p_ma jorancestral_ <* 0.55) were removed.

We used 4-fold and 0-fold degenerate sites as proxies for neutral and selected sites, respectively, and inferred the full DFE (both deleterious and beneficial). The deleterious DFE was parameterized as a gamma distribution with two parameters: *S_d_* (the mean scaled negative selection coefficient) and *b* (the shape parameter). The shape parameter is inversely related to the coefficient of variation, such that higher values of *b* correspond to a less dispersed and less skewed distribution of effects. The beneficial DFE was parameterized as an exponential distribution with two parameters: *p_b_* (the probability that *S >* 0) and *S_b_* (the mean scaled positive selection coefficient). Additionally, we estimated *ε_anc_* for the ancestral allele misidentifica-tion, i.e., polarization error. We compared nested models using likelihood ratio tests (LRTs), assessing whether including additional parameters beyond the deleterious DFE —such as the beneficial DFE or polarization error— significantly improved the model fit. For each inference, we performed 10 independent optimization runs and obtained confidence intervals using 100 bootstrap replicates.

We inferred the DFE both using the full SFS estimated over all genes and using SFSs generated after binning genes into four classes based on their posterior mean selective constraint values from GeneBayes. For each species, genes were assigned to high, moderate–high, moderate–low, and low constraint classes using thresholds of *s >* 1*e^→^*^5^, 1*e^→^*^6^ *< s <* 1*e^→^*^5^, 1*e^→^*^7^ *< s <* 1*e^→^*^6^ and *s <* 1*e^→^*^7^, respectively. These thresholds corresponded to scaled negative selection coefficients (*≈ N_e_s*) of [-Inf, -100), [-100, -10), [-10, -1) and [-1,0) for all species, based on the *N_e_* estimates for these populations (*N_e_ ≈* 5.8 × 10^5^ (Halligan et al., 2010), 1.36 × 10^6^ (Johri et al., 2020), and 9.4 × 10^6^ (Raas and Dutheil, 2024) for mouse, fruit fly and yeast, respectively). We chose these four classes as they represent approximate discrete ranges of selection coefficients from strong to weak purifying selection (Keightley and Eyre-Walker, 2010; Kousathanas and Keightley, 2013; Tataru et al., 2017). To assess the robustness of our results, we also repeated the analyses after dividing genes into 10 and 15 classes based on their posterior mean selective constraint. This time each group included approximately an equal number of 4-fold degenerate polymorphisms. For the DFEs inferred from binned SFS, we compared the likelihood of a joint model (shared parameters across bins) with the product of marginal likelihoods (independent parameters for each class) using LRTs, to evaluate whether parameter sharing was supported by the data. For the joint model, we allowed *ε_anc_* to vary between datasets.

To account for the bias of selecting a single best-fitting model— especially when multiple models had similar likelihoods—we also calculated model-averaged parameter estimates by computing a weighted average across the four DFE models, with weights determined by their respective AIC values (Tataru et al., 2017; Castellano et al., 2019). The SFSs used for DFE inference were also used to compute average heterozygosity at 0-fold (*π*_0_) and 0-fold (*π*_4_) degenerate sites.

Finally, we estimated *α*, the proportion of adaptive substitutions, computed from the beneficial portion of the DFE (equation 10 in Tataru et al. (2017)). We first calculated *α* considering all mutations with *S >* 0 as beneficial. As the inclusion of mutations with very small selection coefficients may cause overestimation, we also calculated *α* after setting a lower bound of 1, thus not considering nearly neutral mutations (Tachida, 1991). We also computed *ω_a_*, the rate of adaptive substitution relative to the neutral substitution rate, as *ω_a_*= *α*(*d*_0_*/d*_4_), where *d*_0_ and *d*_4_ are the per-site divergences at 0-fold and 4-fold degenerate sites, respectively. For the derivation of *ω_a_*, *α* was computed from divergence counts (equation 8 in Tataru et al. (2017)), rather than from the DFE.

## Results

### Informative features about gene-level constraints

We first examined which features were informative for predicting gene-level selective constraints in each species. In all three species, substantial contribution to the prior mean came from conservation scores and gene structure categories (Table 1). The other categories also contributed to varying degrees, and the order was similar between mouse and fruit fly: gene expression had the next strongest contribution, whereas protein–protein interactions contributed the least. For the prior variance, gene structure features were again the largest contributor in yeast, followed by GO terms. In mouse and fruit fly, gene expression contributed the most to the prior variance.

**Table 1:**
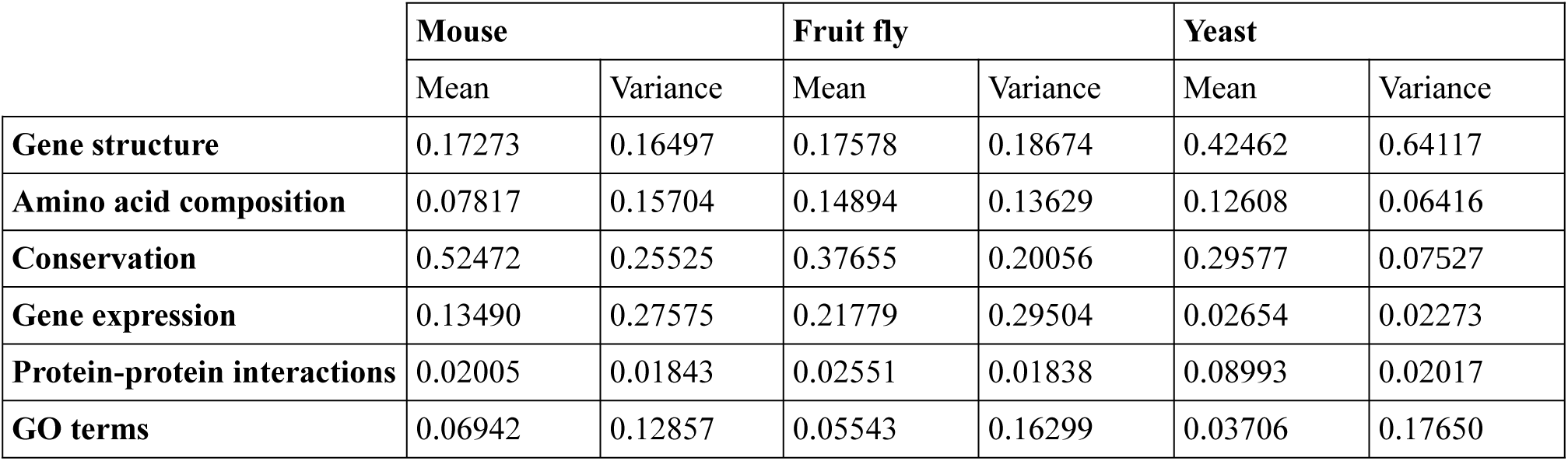
The influence of each gene-feature category on the learned prior mean and variance, shown as the sum of importance metrics for individual features within each category.

|  | **Mouse** |  | **Fruit fly** |  | **Yeast** |  |
| --- | --- | --- | --- | --- | --- | --- |
|  | Mean | Variance | Mean | Variance | Mean | Variance |
| **Gene structure** | 0.17273 | 0.16497 | 0.17578 | 0.18674 | 0.42462 | 0.64117 |
| **Amino acid composition** | 0.07817 | 0.15704 | 0.14894 | 0.13629 | 0.12608 | 0.06416 |
| **Conservation** | 0.52472 | 0.25525 | 0.37655 | 0.20056 | 0.29577 | 0.07527 |
| **Gene expression** | 0.13490 | 0.27575 | 0.21779 | 0.29504 | 0.02654 | 0.02273 |
| **Protein-protein interactions** | 0.02005 | 0.01843 | 0.02551 | 0.01838 | 0.08993 | 0.02017 |
| **GO terms** | 0.06942 | 0.12857 | 0.05543 | 0.16299 | 0.03706 | 0.17650 |

Despite the majority of individual features having an importance score of zero, approximately 7.43%, 11.58%, and 21.82% of all features contributed to some extent to either the prior mean or variance in mouse, fruit fly, and yeast, respectively (Supplementary Figures 1–3). The relative contributions of the individual features within each category were again similar for mouse and fruit fly. For example, within the gene structure category, recombination rate showed the highest contribution to the mean, and CDS length contributed most to the variance in both species. Among GO terms, those with non-zero contributions overlapped between mouse and fruit fly, including regulation of transcription (biological process) and nucleus (cellular component). Further details on feature importance metrics for each species are provided in the Supplementary Data.

The feature importance metrics reported here reflect the relative frequency with which each feature was selected to split the data during model training, weighted by the improvement in the loss function at each split — a standard measure of predictive contribution in gradient-boosted tree models (Duan et al., 2020; Zeng et al., 2024). These values should therefore be interpreted as reflecting the relative contribution of each feature to the learned prior, which reflects their predictive power, rather than the absolute magnitude of their causal effects. Despite this, the consistency of feature contributions across species suggests that the same genomic properties can be broadly predictive of selective constraint.

Additionally, in contrast to linear regression coefficients, gradient-boosted trees select one feature per split sequentially, meaning correlated features compete for splits rather than jointly affecting each other’s contributions. When features are highly correlated, they may substitute for one another across splits, reducing the stability of their individual importance scores without affecting the posterior constraint estimates. Because conservation features are more likely to correlate with many other features, we trained an additional model excluding them (Supplementary Table 4). The resulting classification of genes into the four selective constraint classes based on their posterior estimates (high, moderate–high, moderate–low, and low; see Methods) remained extremely similar across the two models. Based on both hypergeometric and permutation tests, the overlap of genes assigned to each class was significantly higher than expected by chance (Supplementary Table 4).

### Variation in gene features and selective constraints

We next examined how gene features varied with the posterior mean of selective constraints. As a first descriptive summary, we calculated Spearman correlations between individual features and selective constraints. Many features, including those that did not substantially contribute to the prior estimates, exhibited significant correlations (Supplementary Figure 4). Features with absolute Spearman *ρ* values greater than 0.4 belonged to the expression, conservation, or gene structure categories, and values greater than 0.5 were exclusively from conservation features across all species. All features in the expression and PPI categories were positively correlated with selective constraints.

For genes divided into four classes based on posterior estimates of selective constraints, 0-fold and 4-fold diversity (*π*_0_ and *π*_4_) both decreased from the low- to high-constraint classes (Supplementary Figure 5). Similarly, *π*_0_*π*_4_ varied significantly across classes, with the lowest values — corresponding to the highest efficiency of selection — observed in the high-constraint class (Table 2), confirming that this classification captures substantial genome-wide variation.

**Table 2:** π_0_/π_4_ values for whole genome and across selective-constraint classes. The 95% confidence intervals for each point estimate were determined from 1,000 bootstrap resamples and given in brackets.

|  | **Mouse** | **Fruit fly** | **Yeast** |
| --- | --- | --- | --- |
| **Whole genome** | 0.15659 [0.15643-0.15678] | 0.09924 [0.09899-0.09960] | 0.14551 [0.14487- 0.14584] |
| **High constraint** | 0.00519<br>[0.00476-0.00561] | 0.00323<br>[0.00293-0.00356] | 0.00123<br>[0.00058-0.00195] |
| **Moderate-high constraint** | 0.0628<br>[0.0615-0.0639] | 0.0382<br>[0.0374-0.0390] | 0.0500<br>[0.0449-0.0555] |
| **Moderate-low constraint** | 0.231<br>[0.228-0.235] | 0.137<br>[0.135-0.139] | 0.0970<br>[0.0908-0.1044] |
| **Low constraint** | 0.543<br>[0.527-0.561] | 0.390<br>[0.370-0.410] | 0.181<br>[0.176-0.186] |

Across these classes, we examined the distributions of features that showed the highest contributions to the predicted prior or the strongest correlations within each feature category. The distribution of recombination rate and CDS GC content did not show discernible differences between constraint classes for mouse and yeast. In contrast, in fruit fly, CDS GC content showed a slight increase and recombination rate a slight decrease with increasing selective constraint (Figure 2). For amino acid composition features, differences across selective constraint classes were minimal. Conservation scores displayed a clearer relationship with selective constraint in all species: PhastCons values decreased as constraint decreased, while *d_N_/d_S_*and SIFT scores increased as constraint decreased, although this increase was much less pronounced for SIFT (Figure 2; Supplementary Figure 6).

**Figure 2:**
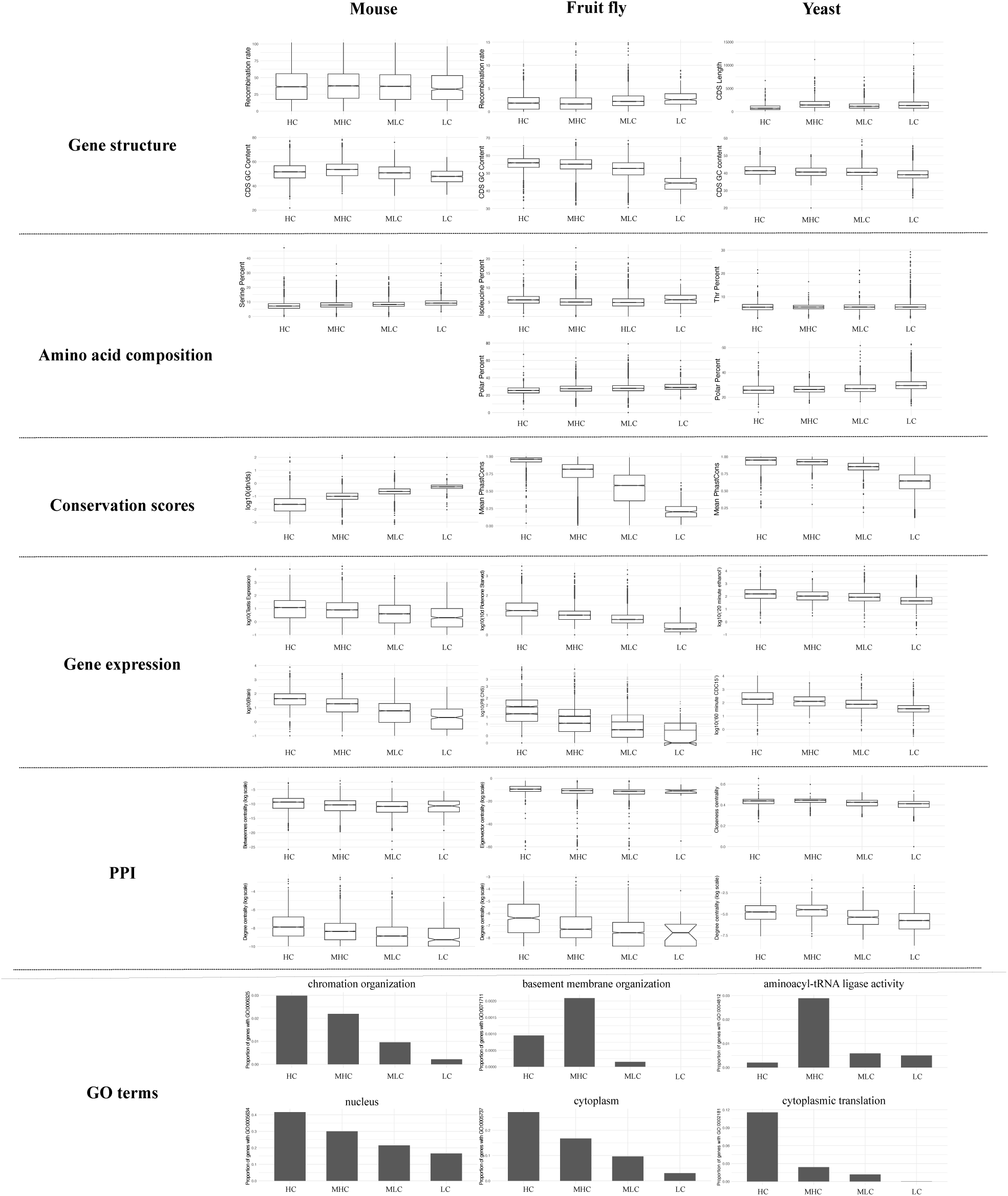
Variation of gene features with selective constraint (HC: High constraint; MHC: Moderate-high constraint; MLC: Moderate-low constraint; LC: Low constraint). For each gene-feature category (rows), the top panel shows the distribution of the feature with the highest contribution to the prior mean, and the bottom panel shows the feature with the strongest Spearman’s ρ correlation. Only a single panel is shown when the top feature is the same for both criteria.

Gene expression features in all species tended to show higher expression levels in more highly constrained genes (Figure 2). Similarly, connectedness measures derived from pro-tein–protein interaction data indicated that genes with greater connectivity and centrality in gene networks were under slightly stronger selective constraint. For non-continuous GO term features, we calculated the proportion of genes annotated with each term within each selective constraint class. GO terms with the highest correlations were associated with nucleus, cytoplasm, and cytoplasmic translation for mouse, fruit fly, and yeast, respectively. In all cases, they showed monotonically decreasing proportions from high- to low-constraint classes. GO terms with larger feature importance metrics for mouse and fruit fly were biological processes related to chromatin and basement membrane organization. For yeast, the top term was aminoacyl-tRNA ligase activity, a molecular function. These terms exhibited higher proportions of genes in the high-constraint or moderate–high constraint classes. In all three species, transcription-and gene expression–related GO terms also contributed substantially to the prior mean, and genes annotated with these terms were more frequently found in the highly constrained classes (Supplementary Figure 7).

Overall, these results indicate that many features are associated with gene-level selective constraint to varying degrees, but no single feature fully explains the observed variation or is sufficient to partition genes, particularly in yeast. Instead, these features are far more predictive of selective pressure across the genome when considered together, in combination. In addition, genes with higher expression levels and greater network connectivity tend to exhibit stronger selective constraint, consistent with previous studies (Hahn and Kern, 2005; He and Zhang, 2006; Mähler et al., 2017; Mack et al., 2019). More importantly, these studies also showed that high expression or a high number of protein–protein interactions are features of highly pleiotropic genes, as such genes are more likely to be present across more tissues and to be involved in more biological processes (He and Zhang, 2006; Barbitoff et al., 2025). Indeed, many studies have used expression (Barbitoff et al., 2025; Guillaume and Otto, 2012) or network connectivity (Josephs et al., 2017; Rennison and Peichel, 2022; Ruelens et al., 2023) as proxies for pleiotropy.

### DFE variation across genome

After inferring per-gene selective constraints and examining the factors contributing to their variation, we next asked whether these constraints correspond to differences in the overall DFE, and whether the DFE varies across the genome. For all species, the marginal DFEs inferred for each constraint class provided a significantly better fit to the data than a joint model (mouse: Δ*L* = 25779.83; fruit fly: Δ*L* = 18636.65; yeast: Δ*L* = 1723.65 and *d f* = 12 for all species), indicating that DFE parameters differ significantly across the genome. The best-fitting models for each constraint class were the same for mouse and fruit fly and only differed for the low-constraint class in yeast (Supplementary Table 5). Highly constrained genes were best described by a purely deleterious DFE. Moderately high– and moderately low–constraint genes were best described by a full DFE (including both deleterious and beneficial mutations). For mouse and fruit fly, the least constrained genes were best modeled by a deleterious-only DFE, whereas in yeast a full DFE provided a better fit. Finally, the best fitting model for the whole genome inference was full DFE in all species.

We first examined the parameters of the deleterious DFE. Across all species, highly constrained genes showed higher mean deleterious selection coefficients, *S_d_*and lower relative variance, corresponding to a higher shape parameter *b* (Figure 3A-B). Analyses dividing genes into 10 (Supplementary Figure 8A) and 15 (Supplementary Figure 9A) constraint classes also revealed stronger deleterious effects for most highly constrained genes. In these finer divisions, genes were grouped by equal numbers of polymorphisms rather than biologically motivated constraint ranges, meaning the bins captured different constraint ranges across species and not necessarily evenly spaced intervals. Therefore, we did not expect to see a clear difference in *S_d_* beyond the highly constrained class. Nevertheless, a decreasing trend in *S_d_* was still apparent, especially for fruit fly. In particular, the increase in the shape parameter *b*- reflecting less dispersion in the distribution of deleterious effects - was clearly observed from low- to high-constraint genes.

**Figure 3:**
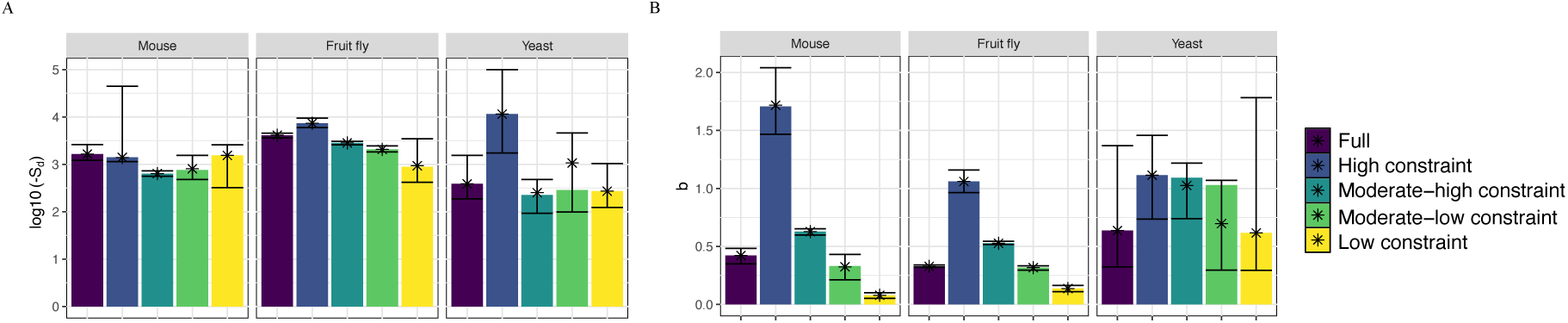
DFE parameters for whole-genome inference and for genes divided into selective-constraint classes. A) Mean deleterious selection coefficient, S_d_. B) The shape parameter of deleterious DFE, *b*. Bars represent parameter values from the best-fitting model, with confidence intervals from bootstrap replicates. Stars indicate the model-averaged parameter estimates.

When comparing whole-genome DFE estimates between species, the variance was smallest for yeast (i.e., highest *b*), and largest for fruit fly. Mean deleterious effect, *S_d_*, was highest for fruit fly and lowest for yeast. Under the usual assumption that organismal complexity increases from yeast to mouse, these between-species patterns did not match the expectations derived from FGM. On the contrary, across-genome variation in the parameters followed the FGM expectations more closely, given that pleiotropy—thus complexity at the gene level—decreased from high- to low-constraint genes, as shown above.

Best fit DFE models for highly constrained genes did not include beneficial mutations in any species. For the remaining classes, mean beneficial selection coefficients (*S_b_*) were higher for the moderate–high constraint class (Supplementary Figure 10A). This pattern did not change when considering the effect of beneficial mutations in every constraint class using model-averaged parameter estimates. Similarly, analyses grouping genes into 10 (Supplementary Figure 8B) and 15 (Supplementary Figure 9B) constraint classes showed that highest *S_b_* values occur in intermediate constraint classes across species. The proportion of beneficial mutations (*p_b_*) tended to increase from high to low constraint (Supplementary Figures 8B-9B-10B). These indicate that, as long as genes are not highly constrained, their beneficial effect sizes can correlate positively with the strength of constraint acting on them, while fewer such mutations are expected.

In comparisons of whole-genome beneficial DFE parameters between species, fruit fly showed the highest *S_b_*, mouse the lowest, and *p_b_* was lower in fruit fly compared to mouse and yeast. Observed *p_b_* values did not align with FGM expectations based on organismal complexity. More complex organisms are expected to have higher proportion of beneficial mutations, as they are expected to be farther away from their fitness optimum (i.e., drift load). However, they are concordant with patterns across species in the efficiency of selection, as measured by *π*_0_/*π*_4_ (Table 2): the efficency of selection was higher for fruit fly, and it was similar between yeast and mouse.

We next examined the proportion of adaptive substitutions, *α*. Highly constrained genes were not inferred to have any adaptive substitutions in any species. Surprisingly, among the remaining classes, we found that *α* decreased from high to low constraint, with the exception of genes in the low-constraint class in yeast (Figure 4, upper panel). With model-averaged parameter estimates, *α* remained highest for the moderate–high constraint class. Additionally, the exception in yeast disappeared when genes were grouped into 10 and 15 constraint classes (Supplementary Figures 11–12); in all species, *α* was higher for genes with intermediate selective constraints and decreased with decreasing constraint. Excluding mutations with low selection coefficients (*S <* 1) did not change this qualitative pattern (Figure 4, Supplementary Figures 11-12, lower panels); only the contribution from lower constraint classes decreased more, as the fraction of mutations with *S >* 1 was lower in their DFEs (Supplementary Figure 13).

**Figure 4:**
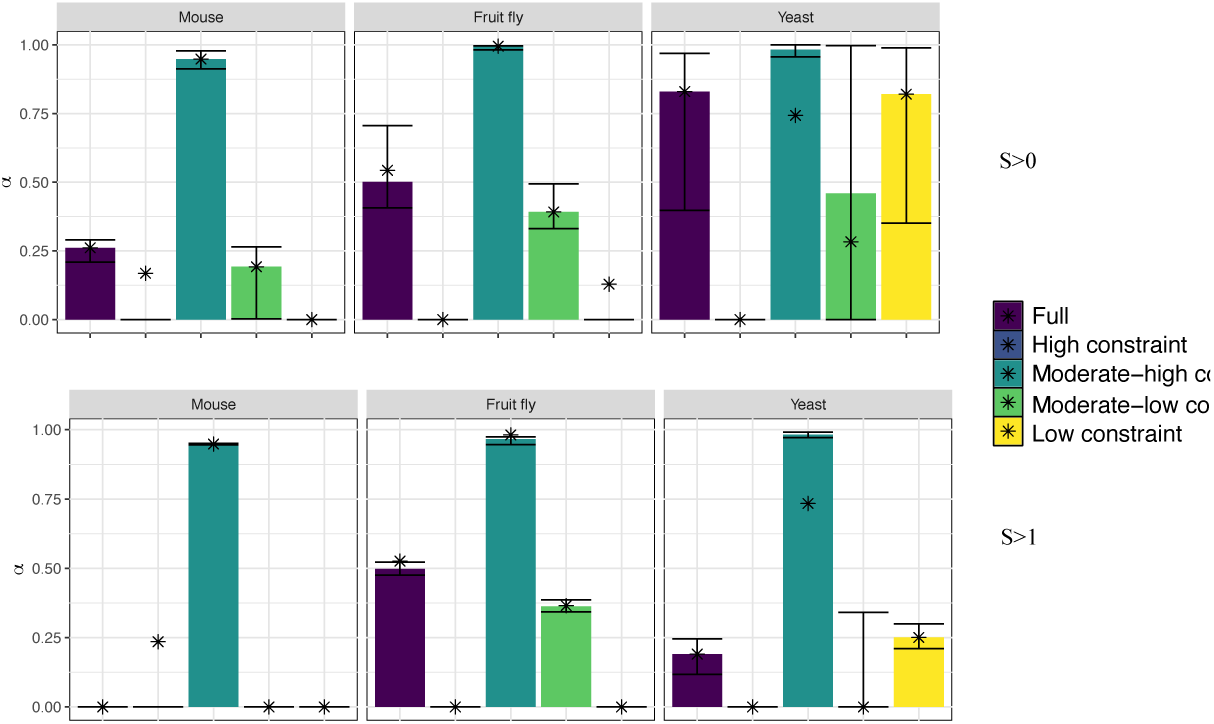
Proportion of adaptive substitutions (α) estimated from the beneficial DFE. Upper panel: including all mutations with S > 0. Lower panel: excluding mutations with low selection coefficients. Bars represent parameter values from the best-fitting model, with confidence intervals from bootstrap replicates. Stars indicate the model-averaged parameter estimates.

As the proportion of adaptive substitutions, *α*, is sensitive to the proportion of nonadaptive substitutions, low *α* in low-constraint bins may reflect the increased rate of nonadaptive substitutions due to reduced efficiency of selection for these low constaint genes. We therefore also computed *ω_a_*, the rate of adaptation. In the four-bin scheme, for mouse and fruit fly the peak estimate of *ω_a_*was in the moderate-low constraint category, whereas the highest estimate of *α* was in the moderate-high constraint category. The confidence interval was wide for the low-constraint bin in both species (Figure 5A). For yeast, the last three bins showed similar values of *ω_a_*. Neither a strictly decreasing nor a clearly concave pattern was consistent across species in this binning scheme. When genes were binned into 10 and 15 constraint classes however, *ω_a_*showed a concave pattern that was consistent across all three species, with higher values at intermediate constraint levels and lower values at both high and low constraint extremes (Figure 5B-C). In contrast to DFE parameters, between-species comparisons of whole-genome *ω_a_* estimates were consistent with FGM expectations based on organismal complexity: *ω_a_* was lower in mouse and fruit fly, and higher in yeast.

**Figure 5:**
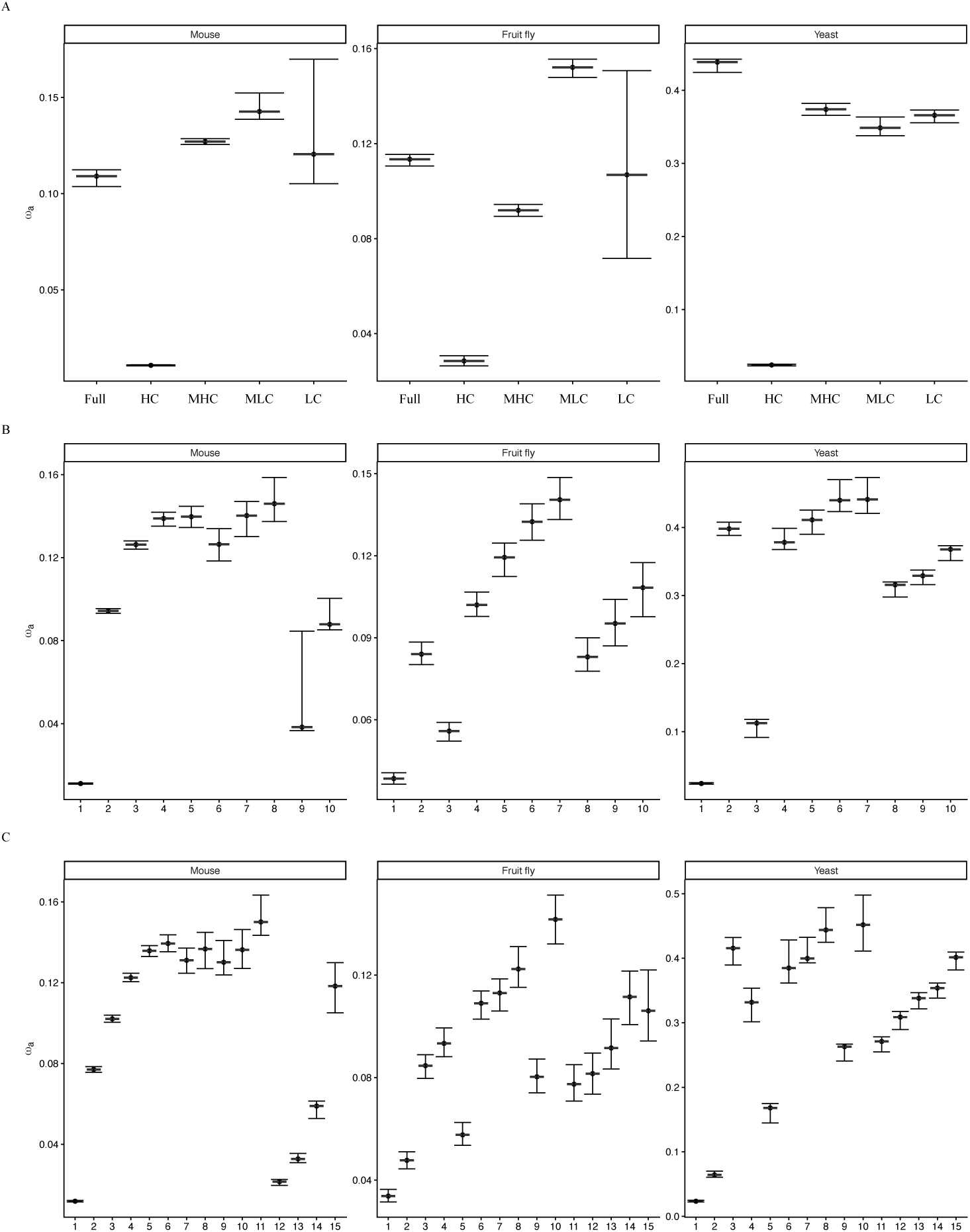
Rate of adaptation (ω_a_) for genes divided into A) four (Full: whole-genome; HC: High constraint; MHC: Modera e-high constraint; MLC: Moderate-low constraint; LC: Low constraint), B) 10, and C) 15 classes based on posterior mean selective constraint estimates. Values were obtained using a lower bound of S>1 (excluding nearly neutral mutations).

Taken together, these results suggest that genome-wide variation in DFE parameters can be explained by differences in complexity at the gene level and follows the predictions of FGM more closely than comparisons based solely on organismal complexity. We therefore asked whether organismal complexity might be reflected in the genome organization of each species, in terms of the relative proportion of genes belonging to each constraint class. There was no pattern that clearly corresponded to organismal complexity, when the four classes examined separately (Figure 6A). Fruit fly and mouse included more genes in moderately-high and moderately-low constrained classes compared to others, which can explain the deviations from FGM expectations in comparisons between species. When the two highly and two weakly constrained classes were combined, a clearer pattern appeared consistent with the complexity of the organisms: the proportion of highly constrained genes was higher for mouse, while the proportion of weakly constrained genes was higher for yeast (Figure 6B).

**Figure 6:**
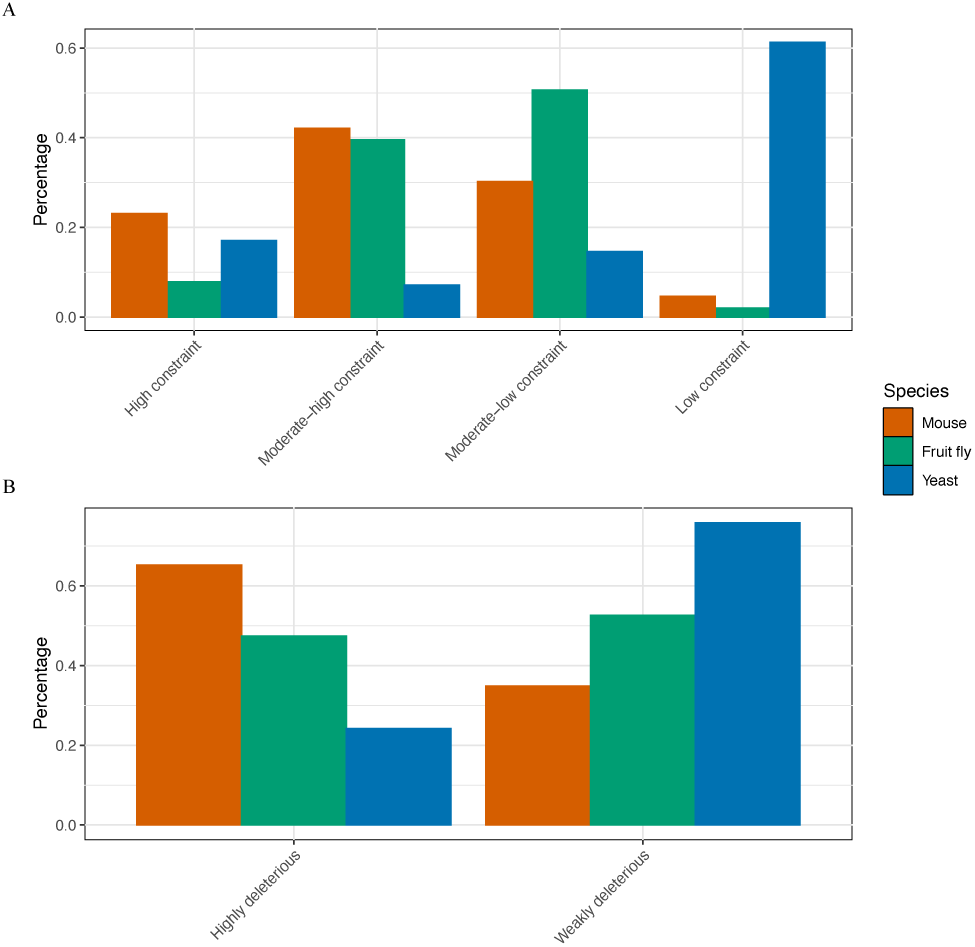
A) The proportion of genes in each selective constraint class. B) Proportion of genes classified as highly deleterious (high + moderate-high constraint) and weakly deleterious (moderate-low + low constraint).

## Discussion

In this study, we examined how gene features predict selective constraints and whether and how the DFE varies across genomes. First, we found that many features affect gene-level constraints. Second, the estimated selective constraints allowed for genes to be categorized into classes with significantly different DFEs. The variation in DFE parameters across genomes follows the predictions of FGM more closely when complexity is considered at the gene level: genes with higher expression and greater connectivity in gene networks — features that can act as proxies for pleiotropy and gene-level complexity — show stronger mean and lower dispersion in deleterious selection coefficients. Between species, the rate of adaptation decreased with increasing organismal complexity, consistent with the cost of complexity theory (Orr, 2000). However, across the genome, the rate of adaptation did not decrease monotonically with gene-level complexity, but tended to be higher at intermediate complexity levels.

Among the three species our results were broadly consistent, with yeast showing departures in a few cases. The order of gene-feature categories based on their contribution to selective constraints was more similar for mouse and fruit fly than for yeast. The most notable difference was the comparatively low contribution of the gene expression category in yeast. This might not solely reflect biological differences between the species, as for yeast there was a lower number of gene expression features available which reduced the variability and predictive signal of this category; the DFE inference for yeast also favored a different best-fitting model for the low-constraint gene class, which may partly reflect the fact that a larger proportion of yeast genes fall under lower selective constraint compared to mouse and fruit fly. When genes were divided into a higher number of classes, this difference disappeared. However, it is important to note that this finer division comes at the cost of evolutionary interpretability: because genes are binned by equal numbers of polymorphisms rather than biologically motivated constraint ranges, individual bins span constraint intervals that are either too narrow or too broad to be meaningfully distinguished. For example, in fruit fly, there are relatively few genes in the low constraint class (Figure 4A), so the lower bins in the finer division end up spanning large constraint ranges that mix nearly neutral and more strongly selected genes together, obscuring the biological signal that the four-class division was designed to capture.

The most informative gene feature categories in predicting selective constraints were, for all species, conservation scores and gene structure. The relationships between constraint and individual features were also largely consistent across the three species. For example, CDS GC content increased with increasing constraint, and higher gene expression and protein–protein interaction were always associated with higher constraint. The same set of categories has also been found to be the best predictors of constraint in humans, with the same directionality of relationships (Zeng et al., 2024), further supporting the generality of these patterns. Not surprisingly, conservation scores were very powerful in explaining selective-constraint variation across the genome. Although the conservation scores we used capture constraint over different evolutionary timescales, they were still related to estimates of deleterious selection coefficients in the current populations. This may indicate that the targets of purifying selection are stable over evolutionary timescales. Additionally, in contrast to *d_N_/d_S_*, the site-based scores Phast-Cons and SIFT are averaged over genes, but this did not reduce resolution, particularly for PhastCons.

Variation in the DFE and other measures of selection with genomic features has been investigated before. These studies typically analyzed genomic predictors separately, aiming to quantify their individual contributions to the observed variation. Our work does not provide such quantification for individual features. Due to the different approaches taken, direct comparisons can be difficult and a one-to-one correspondence cannot be expected. However, our framework still provides information about the relationship between gene features and selective constraints, and our results are consistent with previous findings, especially when compared at macro-scale. In many species, including those examined here, gene expression has been reported as a strong and consistent predictor of purifying selection (Pál et al., 2001b, 2006; Lawrie et al., 2013; Kryuchkova-Mostacci and Robinson-Rechavi, 2015; James and Lascoux, 2025). These studies also showed that gene-structure features such as gene length, GC content, and recombination rate show modest associations with evolutionary constraint, often mediated through covariation with expression (Pál et al., 2001a; Lawrie et al., 2013; Kryuchkova-Mostacci and Robinson-Rechavi, 2015). Similarly, network connectivity has been found to covary with measures of selection (Fraser et al., 2002; Hahn and Kern, 2005; James and Lascoux, 2025). Across species, the degree of variation explained by specific features differs, and there may be species in which associations with certain features are not observed. This may reflect patterns being obscured by noise, or that the marginal effects are truly small. Still, these studies highlight a broadly consistent pattern across taxa: the same classes of genomic features repeatedly emerge as predictors of selective constraint, and our study shows the value of leveraging this biological information to improve gene-level selective constraint estimates.

Investigations of DFE variation across species under FGM expectations have also been conducted, where studies reported support for increasing deleteriousness and decreasing variance in selection strength with increasing complexity (Huber et al., 2017; Lin et al., 2025). However, these comparisons were either limited to *Drosophila* versus human, or summarized as averages at coarse taxonomic levels such as mammals, birds, and insects. The limited resolution and observed deviations may stem from species-specific confounding factors, and also from the difficulty of defining complexity at the organismal level. In contrast, our results for DFE variation across the genome aligned more closely with FGM predictions. Specifically, we found that gene-level complexity — which can be quantified using genomic feature proxies — provides a more informative definition of complexity than organismal labels. We observed that the genomic organization of species — in terms of the relative distribution of genes across constraint classes — was consistent with the expected order of organismal complexity. We therefore suggest that organismal complexity may be better defined from the genomic organization of species, especially when evaluating FGM expectations. Similar arguments about using other measures of complexity, such as genome size or the number of different cell types have been made before (Tenaillon et al., 2007), and our results extend this by suggesting that the gene-level complexity distribution itself can serve as such a measure.

Between species, the rate of adaptation decreased with increasing organismal complexity, considering that from yeast to mouse complexity increases, consistent with the cost of complexity theory (Orr, 2000), and with patterns reported across animal species (Rousselle et al., 2020). However, across the genome, the rate of adaptation did not necessarily decrease monotonically with gene-level complexity, which is not fully consistent with cost of complexity predictions. Some empirical studies have argued that pleiotropy constrains adaptive evolution (Hahn and Kern, 2005; Mähler et al., 2017; Mack et al., 2019). These studies used connectivity and centrality in gene coexpression or protein–protein interaction networks as proxies for pleiotropy and measured selective constraint rather than the rate of adaptation. Since higher pleiotropy correlated with stronger constraint, they concluded reduced adaptive potential. However, this conclusion depends on the assumption that highly constrained genes necessarily adapt less readily. We also observe that highly pleiotropic genes experience stronger selective constraint, but at the same time both the beneficial mutation effect size and the rate of adaptation tend to be higher for genes at intermediate constraint levels.

Similar to our findings, some studies have suggested that varying degree of pleiotropy across genes can result in the highest rates of adaptation occurring in genes with intermediate levels of pleiotropy (Wang et al., 2010; Wagner and Zhang, 2011). Analytical work on the DFE under Fisher’s geometric model, when assuming modular rather than universal pleiotropy, was also in line with our results: the rate of adaptation decreases monotonically with organismal complexity — defined as the total number of dimensions — but shows a concave relationship with gene-level pleiotropy, defined as the number of traits affected per mutation (Lourenço et al., 2011). Empirically, such a nonlinear pattern has been observed in mammals, where highly expressed genes showed higher rates of adaptation except in the highest expression category (Cope et al., 2025). In addition, studies of local adaptation in *A. thaliana* and stickleback demonstrated that intermediate levels of pleiotropy can promote adaptation (Frachon et al., 2017; Rennison and Peichel, 2022). More broadly, numerous other experimental evolution and local adaptation studies have shown that high pleiotropy can contribute to adaptation, without explicitly addressing whether the relationship is monotonic or concave (Hämälä et al., 2020; Thorhölludottir et al., 2023; Whiting et al., 2024; Koch et al., 2025).

We do not argue here that there is no universal pleiotropy. Even if a mutation affects all traits, its pleiotropic effect sizes may vary across traits, especially under a modular organization of gene interactions due to developmental and regulatory processes. Similarly, the omnigenic model of trait evolution proposes that, because of extensive connectivity among gene regulatory networks, most genes can influence most traits, directly or indirectly (Boyle et al., 2017). The model distinguishes between core genes with direct, larger effects, and peripheral genes with indirect, smaller effects. The position of a gene in these networks would define its effective pleiotropy, an observable and quantifiable measure of multifunctionality (Gu, 2014). Indeed, Hill and Zhang (2012) showed that correlations among mutational effects and/or modular organization can yield variable estimates of pleiotropy across genes, even under universal pleiotropy. We argue that the position of genes in networks is relevant, and that effective gene-level measures of complexity and pleiotropyare informative for explaining molecular evolution patterns and DFE variation.

Finally, our finding that within-genome variation in local gene features may better predict variation in the DFE and rate of adaptation than variation among species echoes points made in previous work (Martin, 2014; Soni and Eyre-Walker, 2022; James and Lascoux, 2025). Similarly, the idea that the position of a gene within regulatory networks can define its effective complexity, and that this provides a more useful comparison point within the FGM framework than organismal-level labels, has also been suggested before (Tenaillon, 2014), and is supported by this study. A natural extension of these results would be to ask whether similar variation exists at the within-gene level. For example, since domains are the functional units of proteins and many genes consist of multiple domains, distinct domains within the same gene might show different levels of pleiotropy and therefore different DFEs. Likewise, exons that are differentially included as part of the finished protein through alternative splicing could differ in their pleiotropic effects. Although these questions go beyond the scope of this study, we believe investigating such within-gene variation could be a fruitful direction for future research in order to better understand the scale at which the DFE varies.

## Acknowledgements

We would like to thank Martin Lascoux and Sylvain Glemin for insightful discussions and comments on this work. We also would like to thank Swedish National Infrastructure for Computing (SNIC) for resource allocation for high-power computing and data storage under the project numbers UPPMAX 2025/2-198 and UPPMAX 2026/1-29. This work was supported by the SciLifeLab & Wallenberg Data Driven Life Science Program (grant: KAW 2020.0239)

## Data availability

The data and code used in the analyses, as well as the intermediate data produced, are available in the following repository: https://github.com/Burciny/DFEvariationAcrossGenomes.

