## Supplementary material for "Gene-level complexity explains genome-wide variation in the distribution of fitness effects": Suppl. Fig.

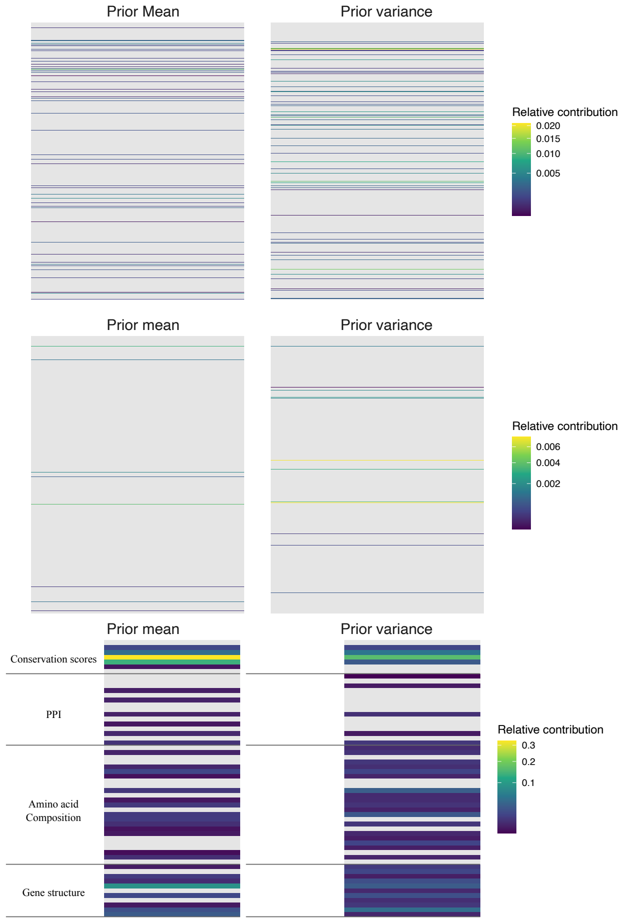


Supplementary Figure 1: Heatmap of the relative contribution of individual features to the prior mean and variance in mouse. The upper panel shows features from the gene expression category, the middle panel shows features from the GO terms category, and the lower panel shows features from all other categories. Grey rows indicate features with an importance score of zero. Underlying values for all features are available at https://github.com/Burciny/DFEvariationAcrossGenomes/GeneBayes_out/Mouse_fullfeature_best_model.feature_importance.tsv

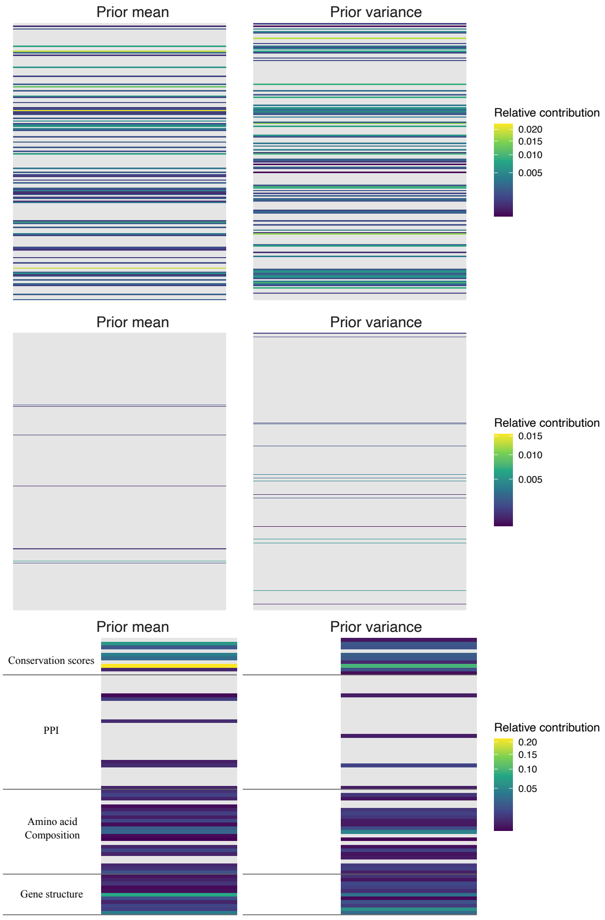


Supplementary Figure 2: Heatmap equivalent to Supplementary Figure 1, showing the results for fruit fly. Underlying values for all features are available at https://github.com/Burciny/DFEvariationAcrossGenomes/GeneBayes_out/Dmel_fullfeature_best_model.feature_importance.tsv

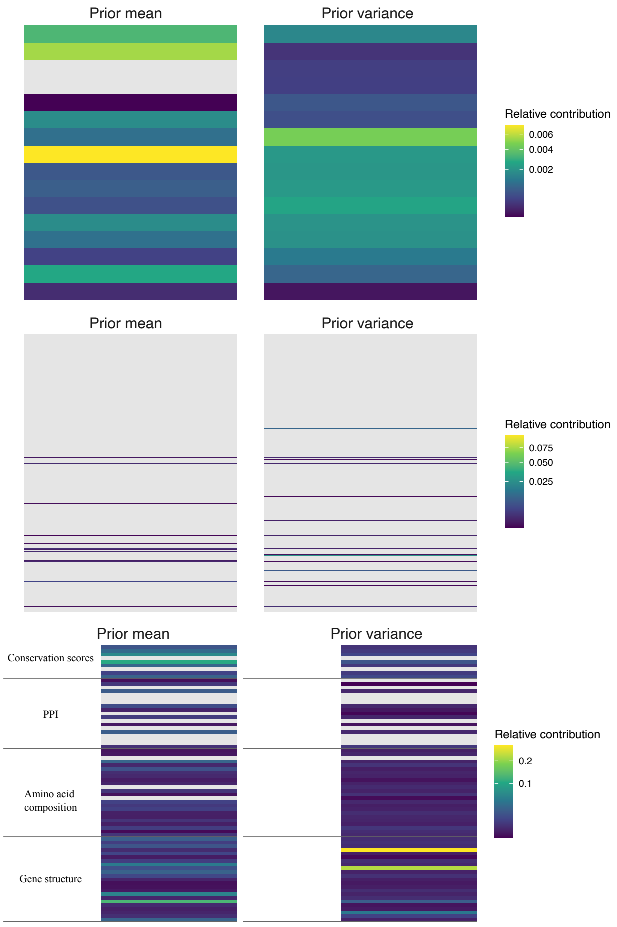


Supplementary Figure 3: Heatmap equivalent to Supplementary Figure 1, showing the results for yeast. Underlying values for all features are available at https://github.com/Burciny/DFEvariationAcrossGenomes/GeneBayes_out/Yeast_fullfeature_best_model.feature_importance.tsv

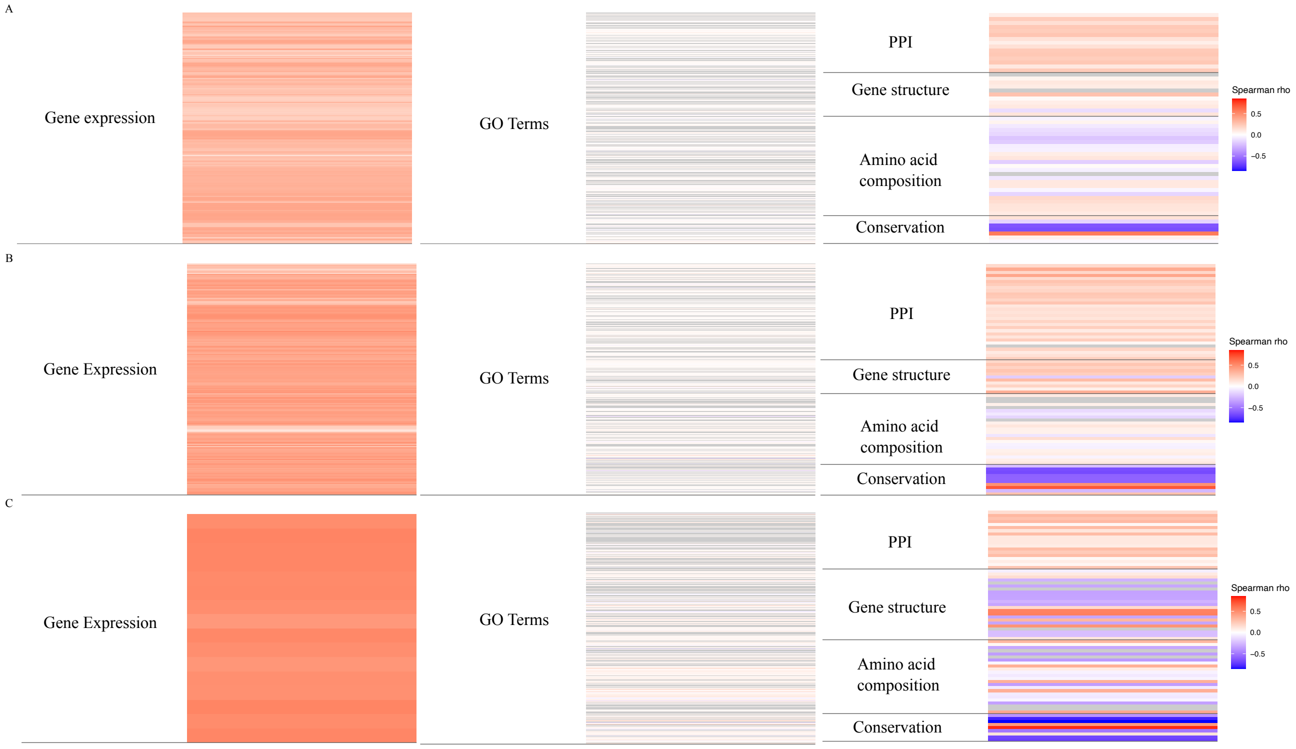


Supplementary Figure 4: Spearman ρ correlations between gene features and per-gene posterior selective constraint for A) mouse, B) fruit fly, and C) yeast. Underlying values for all features are available at https://github.com/Burciny/DFEvariationAcrossGenomes/intermediate_data/postmeanBO_features_cors_{species}

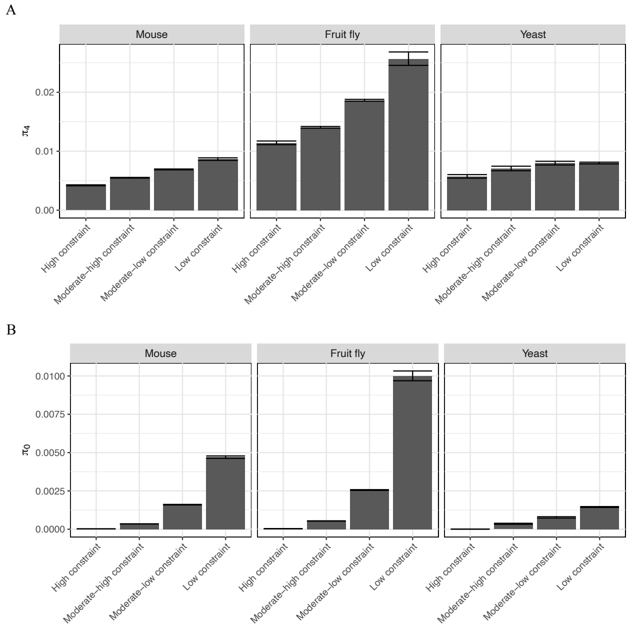


Supplementary Figure 5: A) 4-fold ($\pi_4$) and B) 0-fold degenerate site ($\pi_0$) diversity across selective-constraint classes. The 95% confidence intervals for each point estimate were determined from 1,000 bootstrap resamples and given in brackets.

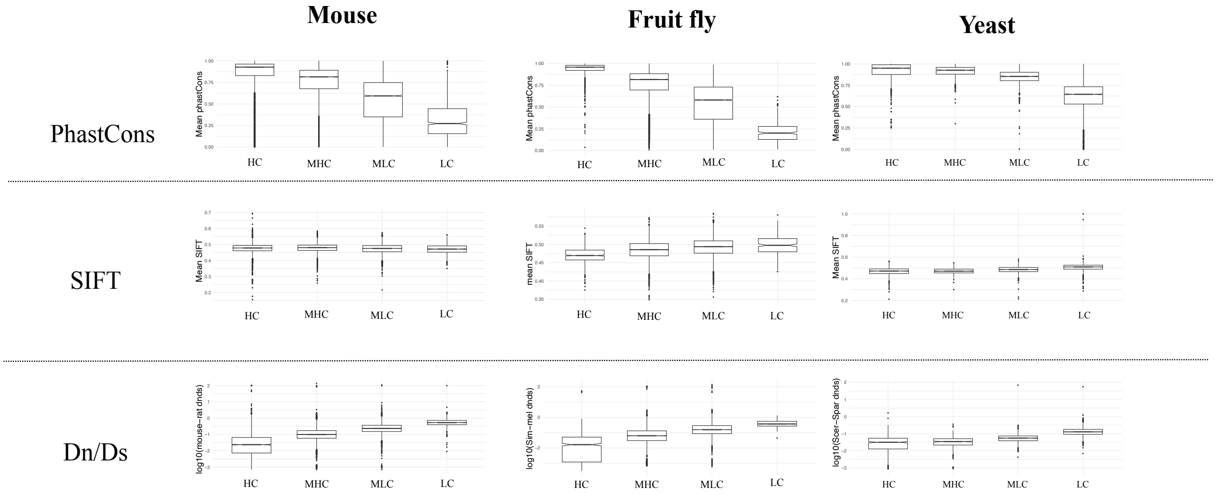


Supplementary Figure 6: Conservation scores across different selective-constraint classes (HC: High constraint; MHC: Moderate-high constraint; MLC: Moderate-low constraint; LC: Low constraint).

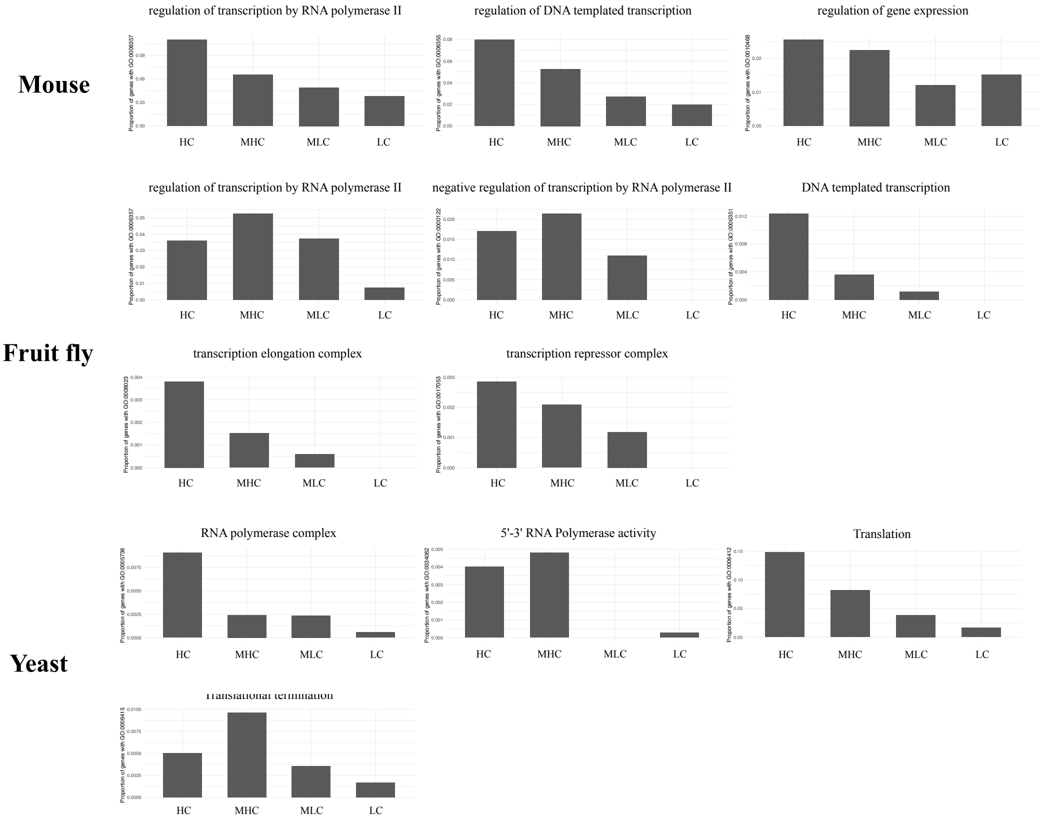


Supplementary Figure 7: Transcription- and gene expression–related GO terms with non-zero contribution to the prior mean in mouse and fruit fly (HC: High constraint; MHC: Moderate-high constraint; MLC: Moderate-low constraint; LC: Low constraint).

A

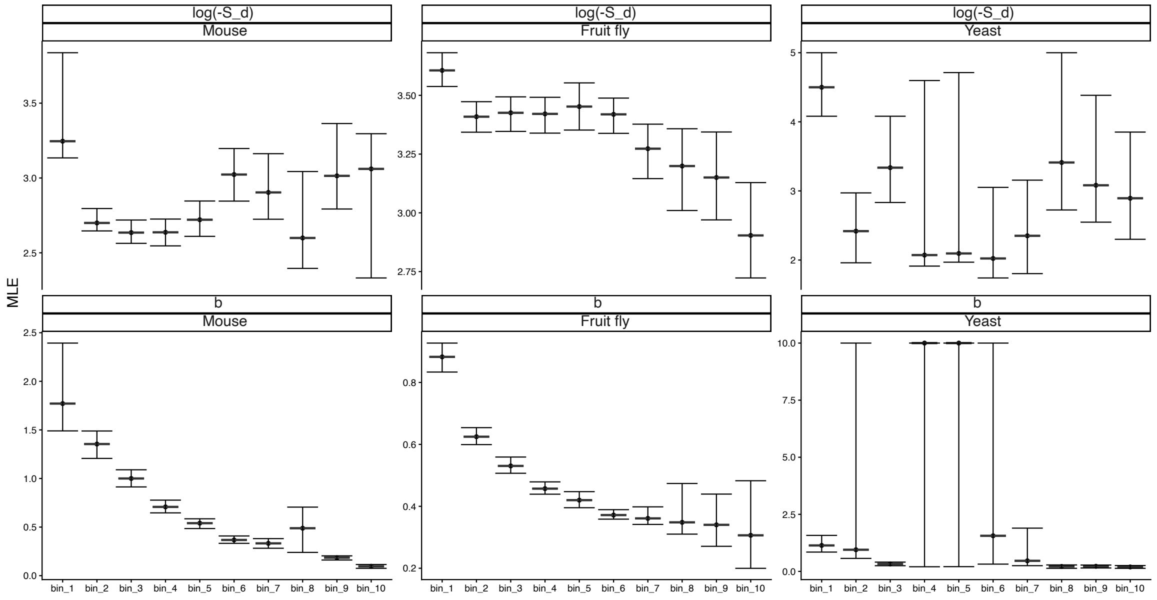


B

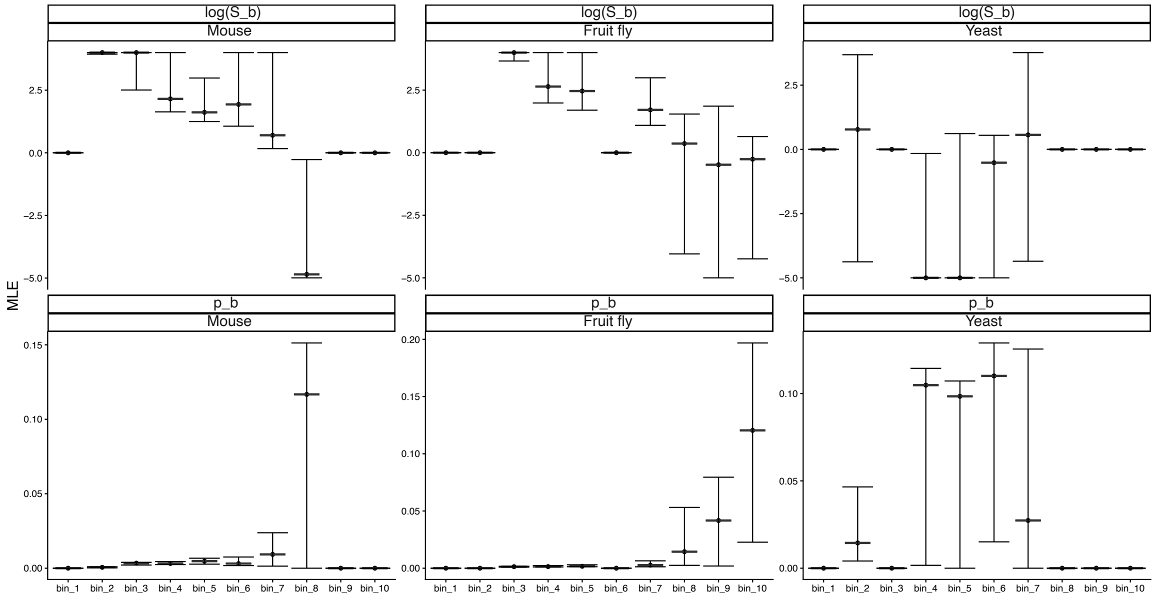


Supplementary Figure 8: DFE parameters for genes divided into 10 classes based on posterior mean selective constraint estimates. Selective constraint decreases from bin1 to bin10. A) Deleterious DFE parameters B) Beneficial DFE parameters C) Rate of adaptation estimated from the beneficial DFE. Upper panel: including all mutations with S > 0. Lower panel: excluding mutations with low selection coefficients

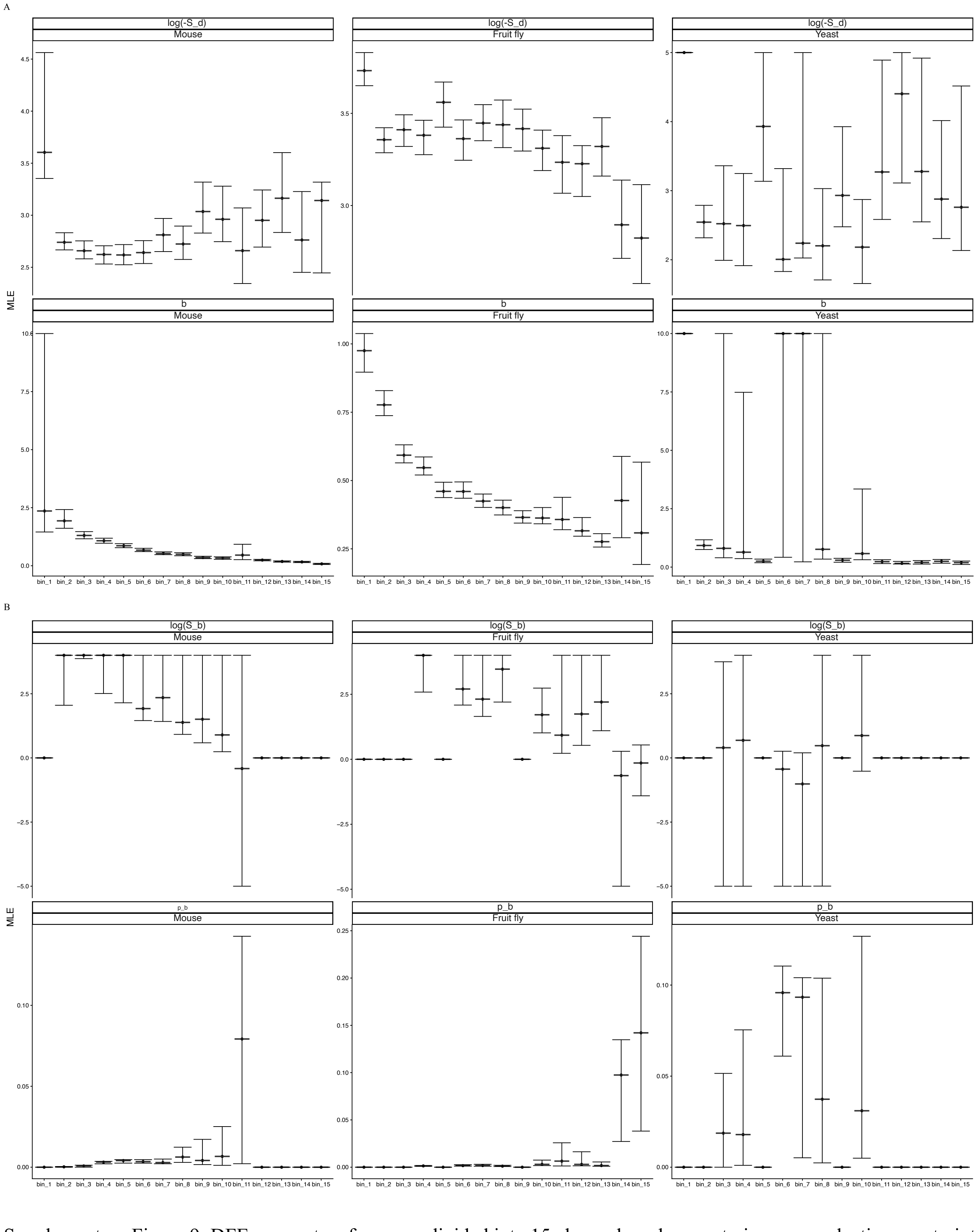


Supplementary Figure 9: DFE parameters for genes divided into 15 classes based on posterior mean selective constraint estimates. Selective constraint decreases from bin1 to bin15. A) Deleterious DFE parameters B) Beneficial DFE parameters

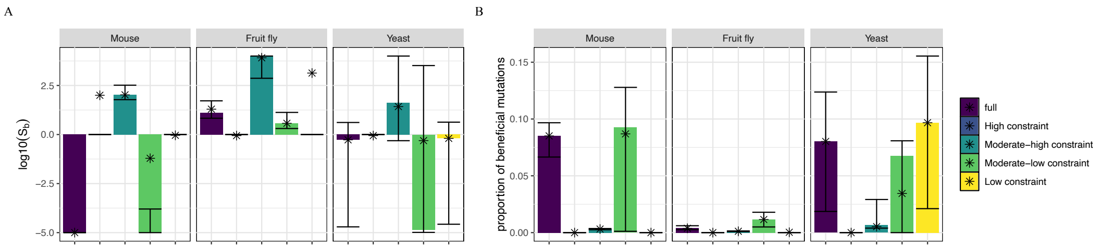


Supplementary Figure 10: DFE parameters for whole-genome inference and for genes divided into selective-constraint classes. A) Mean beneficial selection coefficient, $S_b$. B) Proportion of beneficial mutations, $p_b$. Stars indicate the model-avearaged parameter estimates.

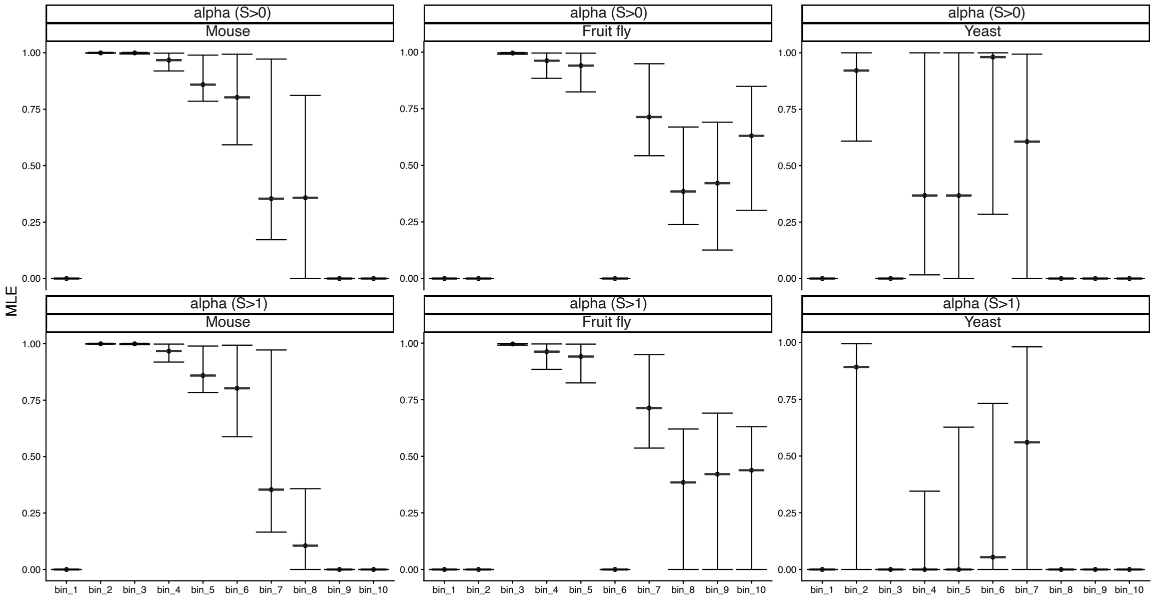


Supplementary Figure 11: Proportion of adaptive substitutions (α) estimated from the beneficial DFE for genes divided into 10 classes based on posterior mean selective constraint estimates. Selective constraint decreases from bin1 to bin10. Upper panel: including all mutations with S > 0. Lower panel: excluding mutations with low selection coefficients (S<1).

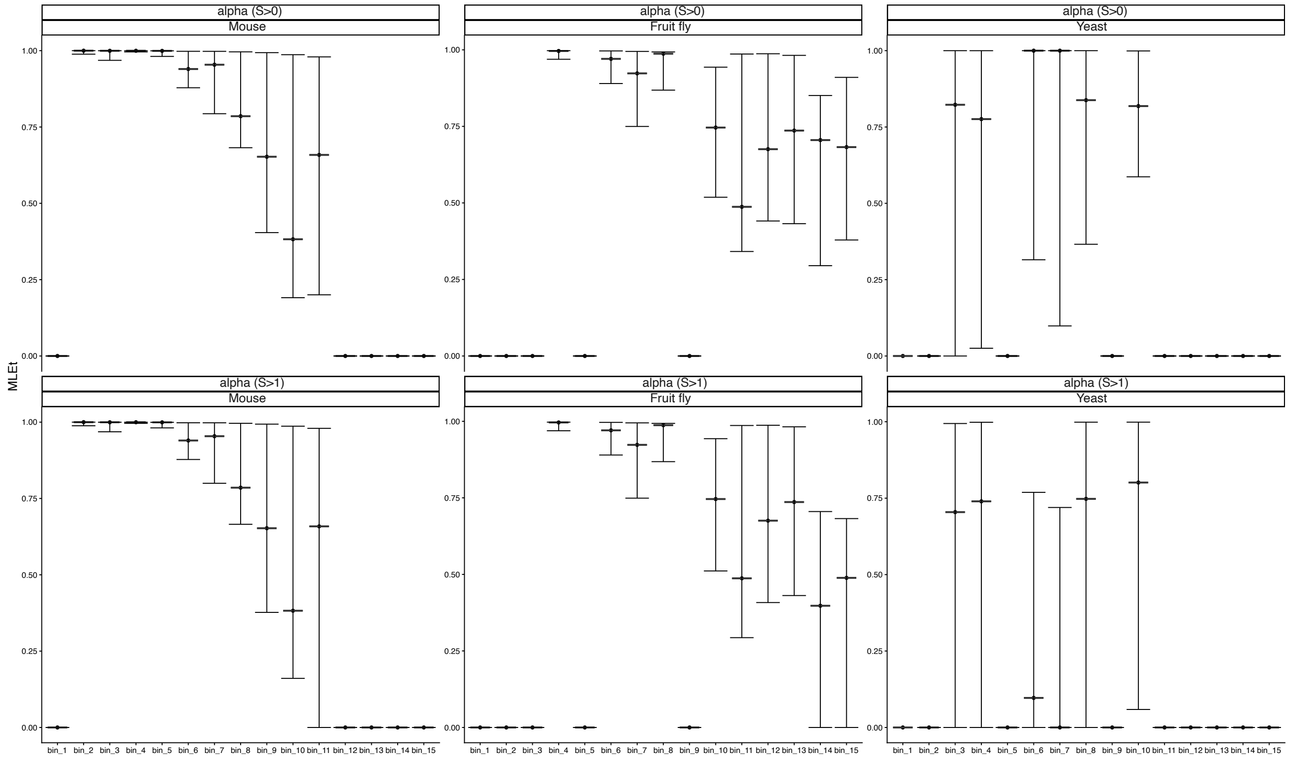


Supplementary Figure 12: Proportion of adaptive substitutions (α) estimated from the beneficial DFE for genes divided into 15 classes based on posterior mean selective constraint estimates. Selective constraint decreases from bin1 to bin15. Upper panel: including all mutations with S > 0. Lower panel: excluding mutations with low selection coefficients (S<1).

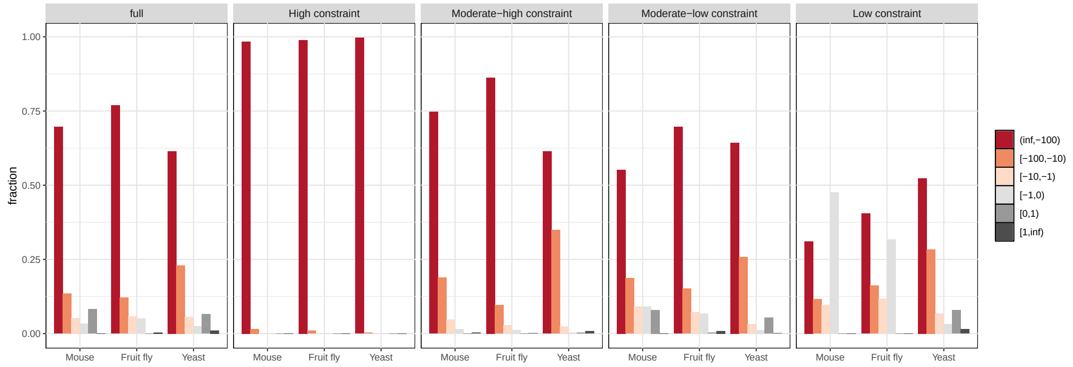


Supplementary Figure 13: Discretized DFE inferred from the whole genome and from genes binned by selective-constraint classes, shown for each species.

Supplementary Table 1a: The number and names of individual gene features in each 6 category before and after filtering for Mouse. Feature names for Gene expression and GO terms can be found in the supplementary data in github repository https://github.com/Burciny/DFEvariationAcrossGenomes/GeneBayes_in/Gene_features_Mouse_[all/filtered].gz

| | All | Filtered |
|---|---|---|
| **Gene structure** | 11 | 11 |
| | - CDS GC content<br>- CDS length<br>- exon count<br>- transcript count<br>- UTR3 length<br>- UTR5 length<br>- UTR3 GC content<br>- UTR5 GC content<br>- transcript length<br>- transcript GC content<br>- recombination rate (cM) | None filtered |
| **Amino acid composition** | 27 | 25 |
| | - 21 aa percent<br>- Total aa<br>- Hydrophilic aa percent<br>- Hydrophobic aa percent<br>- Amphipathic aa percent<br>- Polar aa percent<br>- Nonpolar aa percent<br>- Charged aa percent | Asn and Pro percent filtered out |
| **Conservation** | 9 | 7 |
| | - mean PhastCons<br>- $95^{th}$ percentile PhastCons<br>- max PhastCons<br>- mean SIFT<br>- $95^{th}$ percentile SIFT<br>- max SIFT<br>- dn<br>- ds<br>- dn/ds | Max and $95^{th}$ percentile SIFT filtered out |
| **Gene expression** | 706 | 706 |
| **Protein-protein interactions** | 20 | 15 |
| | - Degree Centrality<br>- Betweennes Centrality<br>- Closeness Centrality<br>- Eigenvector Centrality<br>- Clustering coefficient<br><br>x4 for actual values, decile, quartile, percentile | Degree Centrality decile and quartile; Betweenness Centrality decile and quartile, Clustering coefficient quartile filtered out |
| **GO terms** | 19054 | 3003 |

Supplementary Table 1b: The number and names of individual gene features in each 6 category before and after filtering for Fruit fly. Feature names for Gene expression and GO terms can be found in the supplementary data in github repository https://github.com/Burciny/DFEvariationAcrossGenomes/GeneBayes_in/Gene_features_Dmel_[all/filtered].gz

| | All | Filtered |
|---|---|---|
| **Gene structure** | 11 | 11 |
| | - CDS GC content<br>- CDS length<br>- exon count<br>- transcript count<br>- UTR3 length<br>- UTR5 length<br>- UTR3 GC content<br>- UTR5 GC content<br>- transcript length<br>- transcript GC content<br>- recombination rate (cM) | None filtered |
| **Amino acid composition** | 27 | 23 |
| | - 21 aa percent<br>- Total aa<br>- Hydrophilic aa percent<br>- Hydrophobic aa percent<br>- Amphipathic aa percent<br>- Polar aa percent<br>- Nonpolar aa percent<br>- Charged aa percent | Arg, His, Leu and Lys percent filtered out |
| **Conservation** | 12 | 10 |
| | - mean PhastCons<br>- $95^{th}$ percentile PhastCons<br>- max PhastCons<br>- mean SIFT<br>- $95^{th}$ percentile SIFT<br>- max SIFT<br>- dn x2 (simulans, yakuba)<br>- ds x2 (simulans, yakuba)<br>- dn/ds x2 (simulans, yakuba) | Max and $95^{th}$ percentile SIFT filtered out |
| **Gene expression** | 211 | 211 |
| **Protein-protein interactions** | 20 | 20 |
| | - Degree Centrality<br>- Betweennes Centrality<br>- Closeness Centrality<br>- Eigenvector Centrality<br>- Clustering coefficient<br><br>x4 for actual values, decile, quartile, percentile | None filtered |
| **GO terms** | 9827 | 1735 |

Supplementary Table 1c: The number and names of individual gene features in each 6 category before and after filtering for Yeast. Feature names for Gene expression and GO terms can be found in the supplementary data in github repository https://github.com/Burciny/DFEvariationAcrossGenomes/GeneBayes_in/Gene_features_Yeast_[all/filtered].gz

| | All | Filtered |
|---|---|---|
| **Gene structure** | 25 | 23 |
| | - CDS GC content<br>- CDS length<br>- exon count<br>- UTR3 length<br>- UTR5 length<br>- UTR3 GC content<br>- UTR5 GC content<br>- recombination rate (cM)<br>(Features obtained from SGD https://www.yeastgenome.org/)<br>Mw (molecular weight), PI (isoelectric point), Protein Length, Gravy Score, Aromaticity Score, CAI, Codon Bias, FOP Score, Carbon, Hydrogen, Nitrogen, Oxygen, Sulphur, Instability Index (II), Assuming All Cys Residues Appear As Half Cystines, Assuming No Cys Residues Appear As Half Cystines, Aliphatic Index | Exon count and UTR5 GC content filtered out |
| **Amino acid composition** | 27 | 24 |
| | - 21 aa percent<br>- Total aa<br>- Hydrophilic aa percent<br>- Hydrophobic aa percent<br>- Amphipathic aa percent<br>- Polar aa percent<br>- Nonpolar aa percent<br>- Charged aa percent | Asp, Gln and Tyr percent filtered out |
| **Conservation** | 11 | 9 |
| | - mean PhastCons<br>- 95$^{th}$ percentile PhastCons<br>- max PhastCons<br>- mean SIFT<br>- 95$^{th}$ percentile SIFT<br>- max SIFT<br>- dn<br>- ds<br>- dn/ds<br>- YN00 mean and median dn/ds from 1011 *S. cerevisiae* project (Peter et al. 2018) | Max and 95$^{th}$ percentile SIFT filtered out |
| **Gene expression** | 16 | 16 |
| **Protein-protein interactions** | 20 | 19 |
| | - Degree Centrality<br>- Betweennes Centrality<br>- Closeness Centrality<br>- Eigenvector Centrality<br>- Clustering coefficient<br><br>x4 for actual values, decile, quartile, percentile | Clustering coefficient quartile filtered out |
| **GO terms** | 6002 | 436 |

Supplementary Table 2: Hyperparameters of the models with the lowest validation loss, selected after tuning with grid search and Bayesian optimization. Values are provided for models trained with all six gene-feature categories (full) and without conservation scores (WOcons).

| | **Mouse** | | **Fruit fly** | | **Yeast** | |
|---|---|---|---|---|---|---|
| | Full | WOcons | Full | WOcons | Full | WOcons |
| **lr (Learning rate)** | 0.040 | 0.044 | 0.044 | 0.012 | 0.048 | 0.013 |
| **max_depth** | 2 | 2 | 2 | 2 | 3 | 4 |
| **min_child_weight** | 4.757 | 1.5 | 1.856 | 4.287 | 3.865 | 3.535 |
| **subsample** | 0.888 | 0.7 | 0.905 | 0.838 | 0.770 | 0.909 |
| **n_trees_per_iteration** | 2 | 4 | 2 | 3 | 3 | 2 |
| **reg_alpha** | 3.741 | 4.1 | 1.238 | 3.104 | 4.077 | 3.533 |
| **reg_lambda** | 1.02e-07 | 0.014 | 4.973 | 0.293 | 0.241 | 2.242 |

Supplementary Table 3: Number of genes in each selective-constraint class for GeneBayes models trained with all six categories of gene features (full) and without conservation scores (WOcons). For all species and constraint classes, the number of overlapping genes between the two models was significantly higher than expected based on hypergeometric and permutation tests with 10,000 iterations (p-values $< 1{\times}10^{-6}$).

| | Full | WOcons | Overlap |
|---|---|---|---|
| | **Mouse** | | |
| **High constraint** | 4625 | 5171 | 3872 |
| **Moderate-high constraint** | 8430 | 7929 | 6815 |
| **Moderate-low constraint** | 6054 | 6321 | 5634 |
| **Low constraint** | 921 | 609 | 555 |
| | **Fruit fly** | | |
| **High constraint** | 1050 | 1034 | 894 |
| **Moderate-high constraint** | 5270 | 5466 | 4979 |
| **Moderate-low constraint** | 6754 | 6562 | 6380 |
| **Low constraint** | 265 | 277 | 234 |
| | **Yeast** | | |
| **High constraint** | 994 | 868 | 745 |
| **Moderate-high constraint** | 416 | 297 | 155 |
| **Moderate-low constraint** | 850 | 842 | 358 |
| **Low constraint** | 3574 | 3827 | 3194 |

Supplementary Table 4: Influence of each gene-feature category on the prior mean and variance, excluding conservation-based features. Values are shown as the sum of importance metrics for individual features within each category.

| | **Mouse** | | **Fruit fly** | | **Yeast** | |
|---|---|---|---|---|---|---|
| | Mean | Variance | Mean | Variance | Mean | Variance |
| **Gene structure** | 0.20041 | 0.16587 | 0.28593 | 0.20567 | 0.377 | 0.655 |
| **Amino acid composition** | 0.22231 | 0.10756 | 0.28814 | 0.22081 | 0.037 | 0.049 |
| **Gene expression** | 0.40057 | 0.62808 | 0.33960 | 0.40009 | 0.003 | 0.016 |
| **Protein-protein interactions** | 0.03832 | 0.02892 | 0.03754 | 0.01771 | 0.043 | 0.021 |
| **GO terms** | 0.13838 | 0.06957 | 0.04880 | 0.15571 | 0.538 | 0.258 |

Supplementary Table 5: Best-fit models for each DFE inference, along with their log-likelihood values and number of parameters. Best-fit models for each DFE inference, along with their log-likelihood values and number of parameters. 'Ancestral' and 'no ancestral' refer to models with and without the polarization error parameter ($\varepsilon_{anc}$), respectively.

| | Mouse | Fruit fly | Yeast |
|---|---|---|---|
| Whole genome | **Full** no ancestral<br>nparam=4; L= -188.818 | **Full** ancestral<br>nparam=5; L= -323.685 | **Full** no ancestral<br>nparam=4; L= -34.313 |
| High constraint | **Del.** ancestral<br>nparam=3; L= -17.774 | **Del.** ancestral<br>nparam=3; L= -27.960 | **Del.** ancestral<br>nparam=3; L= -5.251 |
| Moderate-high constraint | **Full** no ancestral<br>nparam=4; L= -168.058 | **Full** no ancestral<br>nparam=4; L= -414.398 | **Full** no ancestral<br>nparam=4; L= -18.822 |
| Moderate-low constraint | **Full** no ancestral<br>nparam=4; L= -109.465 | **Full** ancestral<br>nparam=5; L= -321.407 | **Full** no ancestral<br>nparam=4; L= -24.583 |
| Low constraint | **Del.** no ancestral<br>nparam=2; L= -163.913 | **Del.** ancestral<br>nparam=3; L= -264.662 | **Full** no ancestral<br>nparam=4; L= -35.255 |
